# Natural diversity in the predatory behavior facilitates the establishment of a new robust model strain for nematode-trapping fungi

**DOI:** 10.1101/843698

**Authors:** Ching-Ting Yang, Guillermo Vidal-Diez de Ulzurrun, A. Pedro Gonçalves, Hung-Che Lin, Ching-Wen Chang, Tsung-Yu Huang, Sheng-An Chen, Cheng-Kuo Lai, Isheng J. Tsai, Frank C. Schroeder, Jason E. Stajich, Yen-Ping Hsueh

**Affiliations:** Institute of Molecular Biology, Academia Sinica, 128 Academia Road, Section 2, Nangang, Taipei, Taiwan; Department of Biochemical Science and Technology, National Taiwan University No. 1, Sec. 4, Roosevelt Road, Taipei, Taiwan; Genome and Systems Biology Degree Program, National Taiwan University and Academia Sinica, Taipei, Taiwan; Biodiversity Research Center, Academia Sinica, 128 Academia Road, Section 2, Nangang, Taipei, Taiwan; Boyce Thompson Institute and Department of Chemistry and Chemical Biology, Cornell University, Ithaca, NY 14853, USA; Department of Microbiology and Plant Pathology, University of California, Riverside, 900 University Ave. Riverside, CA 92521, USA

**Keywords:** Nematode-trapping fungi, Predator-prey interaction, Natural population, G protein signaling, Trap morphogenesis.

## Abstract

Nematode-trapping fungi (NTF) are a group of specialized microbial predators that consume nematodes when food sources are limited. Predation is initiated when conserved nematode ascaroside pheromones are sensed, followed by the development of complex trapping devices. To gain insights into the co-evolution of this inter-kingdom predator-prey relationship, we investigated natural populations of nematodes and NTF, that we found to be ubiquitous in soils. *Arthrobotrys* species were sympatric with various nematode species and behaved as generalist predators. The ability to sense prey amongst wild isolates of *A. oligospora* varied greatly, as determined by the number of traps after exposure to *Caenorhabditis elegans*. While some strains were highly sensitive to *C. elegans* and the nematode pheromone ascarosides, others responded only weakly. Furthermore, strains that were highly sensitive to the nematode prey also developed traps faster. The polymorphic nature of trap formation correlated with competency in prey killing, as well as with the phylogeny of *A. oligospora* natural strains, calculated after assembly and annotation of the genomes of twenty isolates. A chromosome level genome assembly and annotation was established for one of the most sensitive wild isolate, and deletion of the only G protein *β* subunit-encoding gene of *A. oligospora* nearly abolished trap formation, implicating G protein signaling in predation. In summary, our study establishes a highly responsive *A. oligospora* wild isolate as a novel model strain for the study of fungal-nematode interactions and demonstrates that trap formation is a fitness character in generalist predators of the NTF family.

**Significance statement:** Nematode-trapping fungi (NTF) are carnivorous microbes that hold potential to be used as biological control agents due to their ability to consume nematodes. In this work we show that NTF are ubiquitous generalist predators found in sympatry with their prey in soil samples. Wild isolates of NTF displayed a naturally diverse ability to execute their predatory lifestyle. We generated a large whole genome sequencing dataset for many of the fungal isolates that will serve as the basis of future projects isolates. In particular, we establish TWF154, a highly responsive strain of *Arthrobotrys oligospora*, as a model strain to study the genetics of NTF. Lastly, we provide evidence that G-protein signaling is necessary for trap induction in NTF.

## Introduction

The living component of soils are in permanent interaction and play central roles in various aspects of biogeochemistry, including nutrient cycling and transport across distances (1). Specifically, nematodes are the most abundant animals in nature, amounting to a staggering 0.3 gigatonnes or 4.5 x 10^20^ individuals, many of which are devastating parasites of plants, animals and humans (2). Although available, nematicide chemicals pose severe environmental and health risks and therefore, biological control agents such as carnivorous predatory fungi hold potential as alternative pest control tools. Predatory fungi that prey and consume nematodes have been described from several major fungal phyla, suggesting that fungal carnivorism has arisen independently multiple times during evolution (3, 4). In particular, nematode-trapping fungi (NTF) constitute a group of nematophagous organisms that includes a large number of species that rely on the formation of traps to ambush their prey (5, 6). We have previously described that NTF eavesdrop on an evolutionarily conserved family of nematode pheromones, the ascarosides, that are used as a signal to switch from saprophytism to predation via the induction of trap morphogenesis (7), and that NTF use olfactory mimicry to lure their prey into traps (8). Upon being trapped, the nematodes are paralyzed, pierced, invaded and digested by the fungus (5, 6). Both nematodes and NTF can be found in a wide range of habitats (9, 10); however, sympatrism and adaptation in the context of their relationships is yet to be investigated and little is known about the natural history of this example of a predator-prey interaction.

In this study, we investigated the ecology of natural populations of nematodes and NTF and found that *Arthrobotrys* species were sympatric with diverse nematodes and behaved as generalist predators. The degree of predation varied greatly amongst strains and several wild isolates were substantially more efficient predators than the currently used laboratory strain. We sequenced and assembled the genomes of a number of wild *A. oligospora* strains and demonstrated a correlation between the levels of trap formation, prey killing efficiency and phylogeny. Lastly, targeted gene deletion of the G protein *β* subunit-encoding gene revealed that the predatory lifestyle of *A. oligospora* is dependent upon G protein signaling.

## Results and Discussion

### Soil samples reveal a rich diversity of nematodes and NTF

With the goal of isolating sympatric nematodes and NTF from natural habitats, we collected 178 soil samples from 69 ecologically diverse sites in Taiwan (Fig. 1A). Sampling sites were mainly located in northern and central Taiwan, and included forest, lake, agricultural, and coastal environments, as well as educational campuses. More than 66% of the sampling sites were mountainous areas, with altitudes ranging from ∼300-3400 m (Table S2). We established pure cultures of both organisms and their identities were confirmed by sequencing the internal transcribed spacer (ITS) region of the fungal strains and the small subunit (SSU) rDNA regions of the nematodes. By not applying a metagenomic sequencing approach, we have likely underestimated the fungal and nematode biodiversity in our samples. Nevertheless, our approach enabled us to establish cultivable isolates of NTF and nematodes, and examine their interactions in the laboratory. We identified nematodes and NTF in the vast majority of the 178 soil samples and ∼92% and ∼68% of the sampling sites contained nematodes and/or NTF, respectively. They were sympatric in more than 63% of the sites, indicating that these organisms share the same niches in soil environments and interact closely in nature (Fig. 1B). Only two sampling sites (the Xiaoyoukeng fumarole at Yangmingshan National Park and the coastline of Penghu Island) were found to be free of nematodes and NTF, presumably due to extreme environments (high temperature and high salt concentration, respectively) that likely impede survival. In total, we identified 16 previously described species of NTF, as well as a few strains that could represent new species. Among the 16 known species, 11 were identified as belonging to the genus *Arthrobotrys*, which develops three-dimensional adhesive nets as a trapping device to capture prey (5, 6) (Fig. 1C). *A. oligospora* was the most commonly found species, followed by *A. thaumasia* and *A. musiformis*; combined, these three species were present in ∼70% of the sampling sites that had been identified as harboring NTF (Fig. 1C). The remaining five species were of the *Monacrosporium* and *Drechslerella* genera, which develop adhesive columns and constricting rings as trapping devices (11), and were each isolated from one sampling site only (Fig. 1C). Approximately 35% of the sampling sites contained two or more species of NTF, indicating that different species of NTF can occupy the same ecological niches (Table S2). Conidial morphology was diverse amongst the isolated NTF (Fig. 1C).

**Figure 1.**
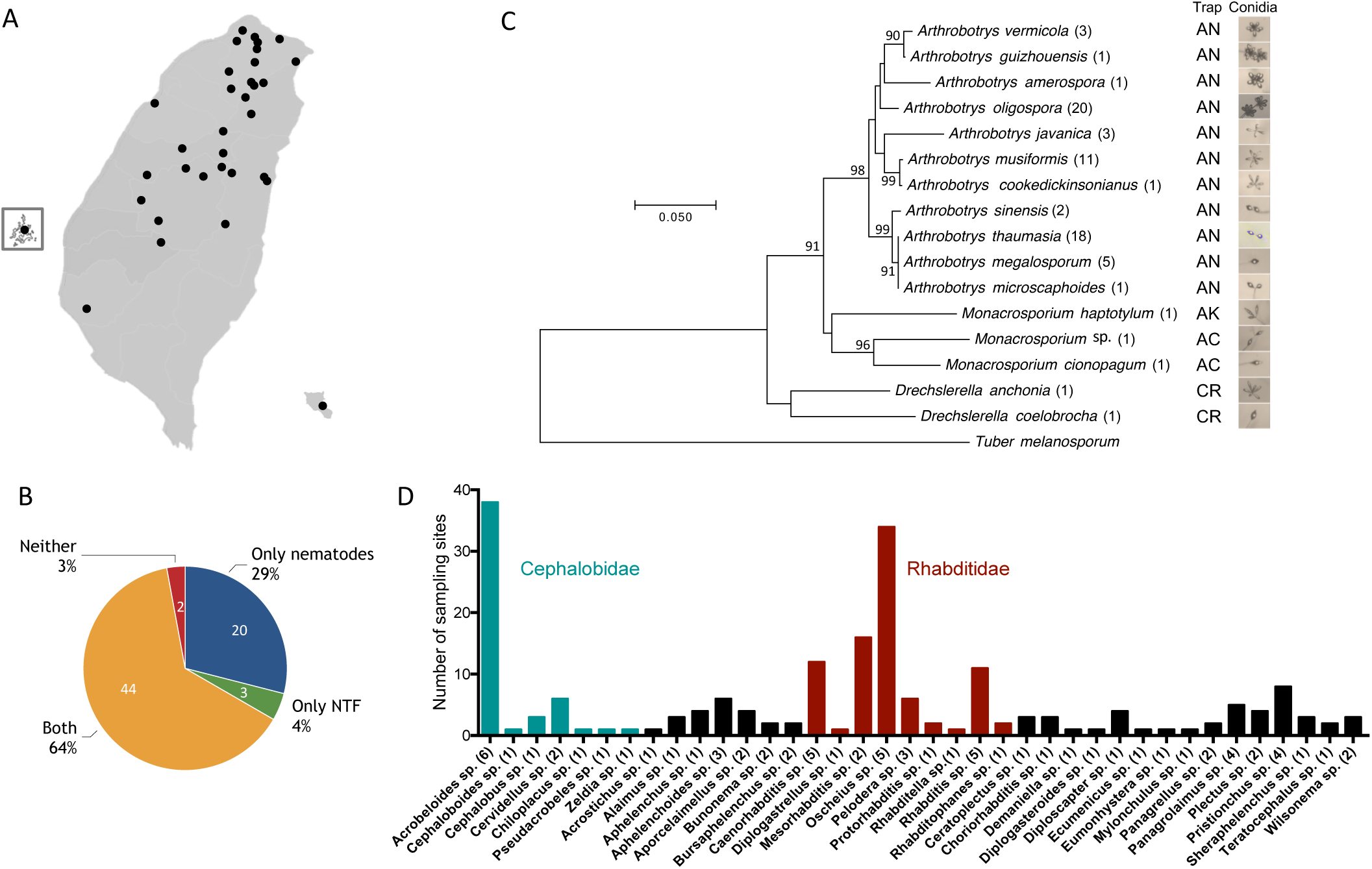
High diversity of nematodes and NTF species are sympatric in soils samples. (A) Geographic distribution of sampling sites in Taiwan. (B) Nematodes and NTF are sympatric in more than 60% of our sampling sites. (C) Phylogeny of the different species of NTF isolated in this study based on their ITS sequences. The numbers represent the prevalence of the difference species of NTF in soil samples. The trap types and images of conidia are shown at the right. AN: adhesive networks, AK: adhesive knobs, AC: adhesive columns, CR: constricting rings. (D) Prevalence and diversity of nematode species in Taiwanese soil samples.

Nematodes were present in ∼93% of the soil samples (Fig. 1B). We cultured the wild nematodes on standard *C. elegans* NGM medium with *Escherichia coli* (OP50) as food source, and directly amplified and sequenced the SSU rDNA regions from those able to grow on OP50. We identified more than 74 Nematoda species (Fig. 1D), covering clades II, IV and V (Fig. S1), demonstrating a rich nematode biodiversity in Taiwanese soil environments. *Oscheius tipulae, Acrobeloides apiculatus,* and *Acrobeloides nanus* were found in ∼40%, ∼30%, and ∼9% of the sampling sites, respectively (Fig. 1D), in line with previous surveys that showed that these nematodes, especially *Oscheius tipulae,* are widespread cosmopolitan soil nematodes (12–16).

### *Arthrobotrys* fungi behave as generalist predators

*A. oligospora*, *A. thaumusia* and *A. musiformis,* the three most commonly identified NTF in our soil samples, were sympatric with at least 50, 29, and 31 species of nematodes, respectively, indicating that NTF naturally encounter a broad range of nematodes, including several *Caenorhabditis* species (Table S3). *Caenorhabditis elegans* is known to be associated with rotten fruits (17), and in one soil sample collected from an apple orchard, we identified both *C. elegans* and *A. oligospora*, providing direct evidence that *C. elegans* encounters *A. oligospora* in its natural habitat. To assess nematode predation, we tested the trapping ability of *A. oligospora*, *A. thaumasia* and *A. musiformis* on sympatric nematode species and observed that all three *Arthrobotrys* preyed on all of the nematode species for which we were able to establish cultures (Fig. 2A). Importantly, when nematodes and NTF from distant sampling sites were co-cultured, trap formation and carnivorism was still observed. For example, the nematodes *Pristionchus pacificus* and *Cervidellus vexilliger* were not sympatric with *A. oligospora* in any of the samples, yet *A. oligospora* still developed traps and consumed isolates of these two nematode species (Fig. 2B). Thus, we conclude that *A. oligospora* does not specifically recognize and prey on particular species of nematodes, but instead behaves as a generalist predator. These data support previous finding that highly conserved nematode signals such as ascarosides play a critical role in this predator-prey interaction (7). Additionally, our observation that multiple NTF species co-exist in the same sample suggests that they might compete for prey in their natural environments.

**Figure 2.**
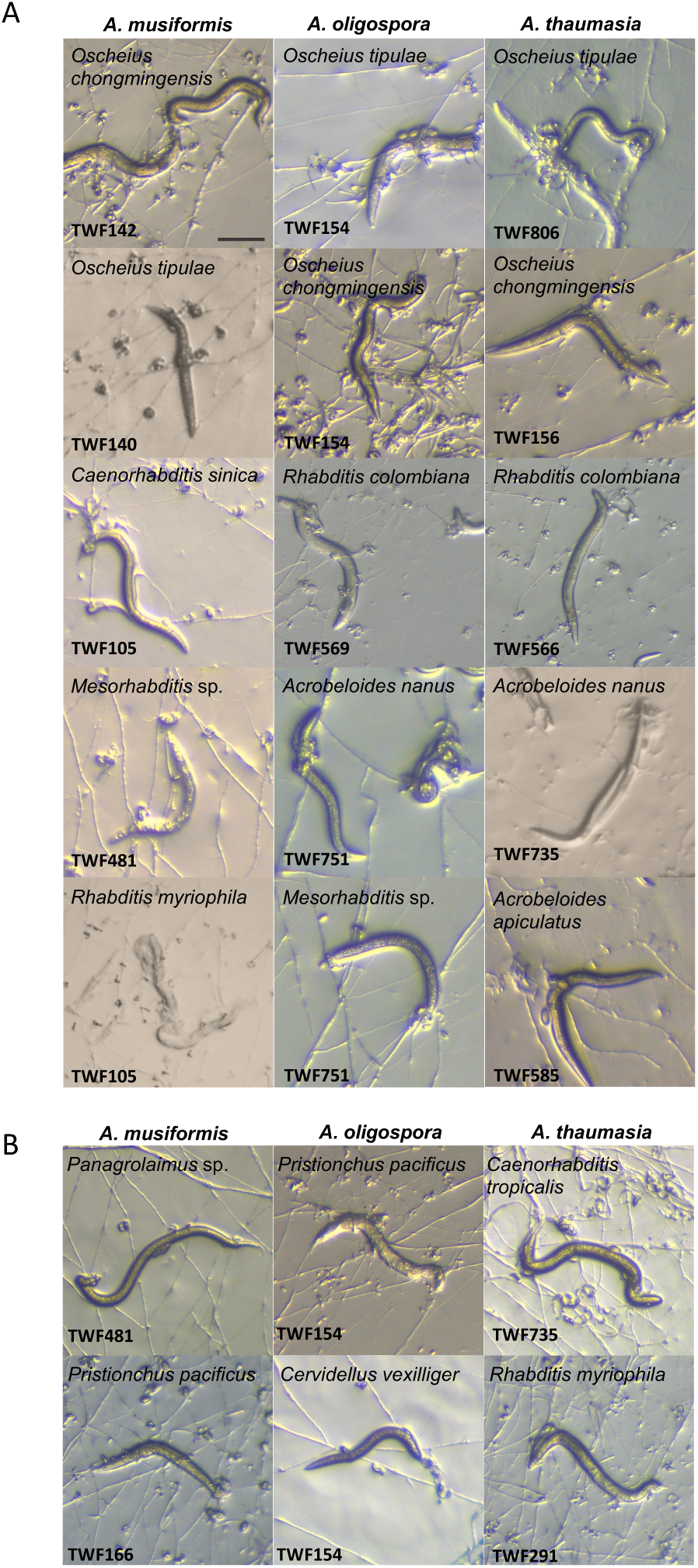
*A. oligospora*, *A. musiformis* and *A. thaumasia* are generalist predators. (A) NTF preying on different sympatric nematode species. Bar = 200 µm (B) NTF preying on allopatric nematode species.

### Predation is a highly polymorphic trait in *Arthrobotrys* species

Phenotypic diversity is commonly observed in natural populations of fungi. For example, variations in stress sensitivity have been observed in *Saccharomyces cerevisiae* (18) and *Neurospora crassa* (19). Two previous studies have surveyed *A. oligospora* in the wild (20, 21) but no phenotypical characterization has been performed. Thus, we designed an assay to quantitatively measure the ability of individual isolates to sense prey and develop traps, the most distinctive feature of NTF, and examined the natural diversity of nematode-trapping performance. Wild NTF were exposed to 30 *C. elegans* L4 larvae for 6 hours and the number of traps developed in cultures at 24 hours post-exposure to nematodes was determined. The average prey-sensing ability of 35 of the *A. oligospora* wild isolates was highly polymorphic for this particular trait, ranging from ∼8 to 162 traps (Fig. 3A). Noteworthy, even TWF788, the lowest performing isolate from our samples, formed, on average, a higher number of traps than the most commonly studied laboratory strain of *A. oligospora*, ATCC24927 (22), while the highest performing strains, TWF132 and TWF154 (Fig. S2), exhibited almost 20-fold more traps than ATCC24927. In fact, our quantitative approach revealed several statistically different groups and at least 21 wild isolates outperformed ATCC24927. A polymorphic response that ranged from ∼8 to 92 traps was also observed when fungal hyphae were exposed to synthesized ascaroside #18. While some wild isolates were highly responsive, others were almost insensitive to this cue, and the ATCC24927 was again amongst the lowest performing statistical group (Fig. 3B). This finding demonstrates that ascaroside-triggered trap morphogenesis is highly polymorphic among our *A. oligospora* wild isolates. Responsiveness to *C. elegans* was positively correlated with responsiveness to ascarosides (Fig. S3). This correlation, despite robust (Spearman, r=0.7556, *p*-value < 0.0001), was not observed throughout all the strains suggesting that prey sensing is a multifactorial process. We did not observe a clear pattern when the association between geographic location and prey-sensing phenotypes was examined (Fig. S4). We further determined trap formation in 10 *A. thaumusia* and 10 *A. musiformis* wild isolates in response to *C. elegans* (Fig. S5), and found that this trait was also highly polymorphic in these two species. Thus, providing evidence that prey sensing is likely highly polymorphic in natural populations of the *Arthrobotrys* genus and that *A. thaumasia* strains formed substantially less traps than *A. oligospora* and *A. musiformis* (Fig. S5). We next investigated if the polymorphic prey-sensing phenotype displayed by *A. oligospora* wild isolates was nematode species-dependent. We selected three *A. oligospora* strains that were strongly responsive (TWF132, TWF154, TWF145) and four strains that were weakly responsive (TWF106, TWF788, TWF191, ATCC24927) to *C. elegans* and measured their ability to form traps in response to four other nematode species (*C. briggsae*, *O. tipulae*, *Rhabditis rainai* and *P. pacificus*). Overall, the levels of trap formation were diverse amongst these strains and if responsiveness to *C. elegans* was higher, the sensitivity to other nematode species was similarly higher and *vice versa* (Fig.3C). Thus, natural diversity in the quantities of trap formation is not restricted to just a single species of NTF nor it is restricted to the response to a single species of nematodes.

**Figure. 3.**
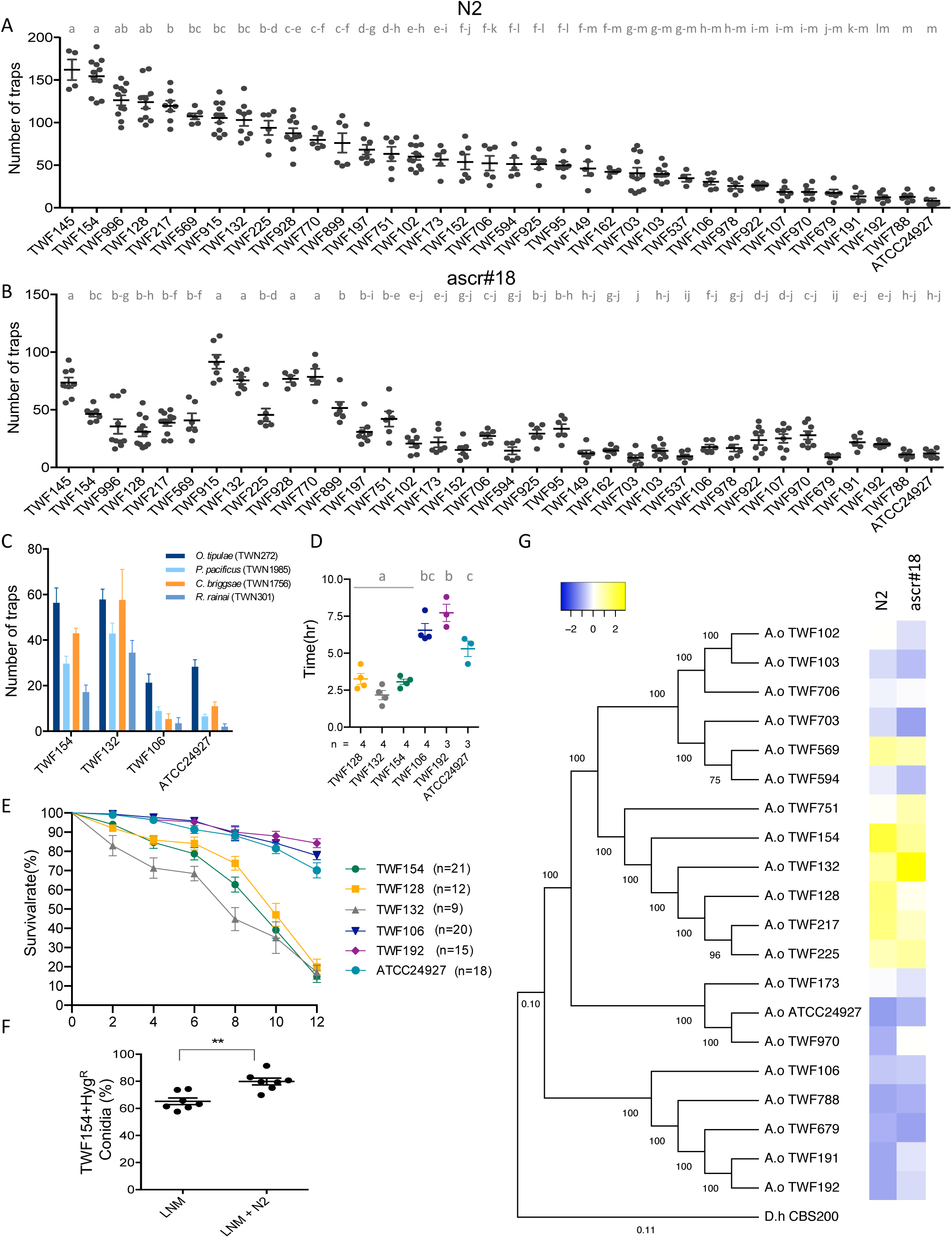
Prey-sensing ability varies considerably among wild isolates of *A. oligospora*. (A) Quantification of trap number induced by *C. elegans* wild-type strain N2 among wild isolates of *A. oligospora* (mean ± SEM, n=6). (B) Quantification of trap number induced by ascarosides among wild isolates of *A. oligospora* (mean ± SEM, n=6). (C) Quantification of trap number induced by *C. elegans* in wild isolates of *A. musiformis* and *A. thaumasia* (mean ± SEM, n=6). (D) Time of trap emergence after exposition to *C. elegans* among different *A. oligospora* wild isolates. (E) Percentage of alive *C. elegans* after exposition to different strains of *A. oligospora* over 12 hours. (F) Competition fitness assay of *A. oligospora* isolates TWF154 and TWF106, computed as the percentage of conidia produced by TWF154 against the total number of conidia, in the presence and absence of *C. elegans* (N2). (G) Phylogeny of 19 *A. oligospora* wild isolates sequenced and assembled for this study. Phylogenetic tree was constructed using 500 random single orthologs and the heatmap summarizes the trapping response of the isolates towards N2 and ascarosides.

We next assessed whether the onset of trap morphogenesis and the ability to kill nematode prey varied among the natural population of *A. oligospora* in addition to the variation in trap numbers developed in response to nematode prey. We found that strains TWF154, TWF128 and TWF132 which formed more traps in response to nematode prey (Fig. 3A), developed the initial traps as soon as ∼2.5 hours after exposure to *C. elegans* whereas strains forming fewer traps (TWF106, TWF192 and ATCC24927) exhibited slower onset of trap morphogenesis (∼5 to ∼7.5 hours) (Fig. 3D). In line with these results, strains that developed traps faster and greater in number captured the nematode prey significantly faster (Fig. 3E). In our killing assays, only ∼20% of nematodes were still alive 12 hours after being added to the fungal cultures in the presence of highly responsive wild isolates in contrast to ∼75-85% of survival in the case of the weakly responsive wild isolates (Fig. 3E). Altogether, these results indicate that predation on nematodes is a naturally diverse trait in populations of generalist NTF and that some strains have evolved to become intrinsically more robust and efficient in sensing nematodes, executing the trapping developmental program and consuming prey. Hence, trap morphogenesis seems to function as a fitness character in NTF.

### Competition assays link predation ability to fitness in *A. oligospora*

After consuming the nematode prey, *A. oligospora* develops chains of conidia (asexual spores) (Fig. S6). To test the hypothesis that predation ability contributes to the fitness of *A. oligospora*, measured by the number of conidia produced, we genetically marked the strong predator TWF154 with a hygromycin-resistant marker (TWF154-*HYG*^R^), and competed this strain with the weak predator TWF106 in the presence or absence of nematode prey. We found that in the absence of nematode prey, the strong predatory strain exhibited a moderate fitness advantage than the weak predatory strain; ∼62% of the conidia harvested from the competition assay were hygromycin resistant (Fig. 3F). In contrast, in the presence of nematode prey, the fitness advantage was further increased by ∼15%; since 78% of the conidia harvested from the competition assay were produced by the strong predatory strain (Fig. 3F). These results suggest that higher predation ability results in better fitness in the nematode-trapping fungus *A. oligospora*.

### The nuclear and mitochondrial genome of a highly responsive *A. oligospora* isolate

In order to generate a genomic resource for future studies on the molecular basis of trap morphogenesis in NTF, we assembled the genome of 19 *A. oligospora* wild isolates (Fig 3G). Interestingly, a phylogenetic analysis using 500 random genes from these 19 genomes (and ATCC24927) showed a clustering pattern that was correlated with the levels of trap formation (Fig. 3G). For example, TWF751, TWF154, TWF132, TWF128, TWF217 and TWF225 formed a separate clade and belonged to the highest performance group which were most sensitive to the nematode prey (Fig. 3A). We anticipate that future genome-wide association studies and comparative genomics analyses with additional genomes from the natural population will disclose multiple candidate genetic loci that govern the predation trait.

We decided to focus on TWF154, a high-performance wild isolate, both in response to *C. elegans*, other nematode species and ascarosides (Fig. 3A, B and C), that we intend to establish as the preferred *A. oligospora* model wild-type strain for further molecular mechanistic studies. We used the PacBio RSII platform to obtain high quality long reads from strain TWF154 with 131X coverage and assembled a complete and contiguous genome of 39.6 Mbp distributed over 10 scaffolds (Fig. 4A). Eight out of ten contigs represented a telomere to telomere complete chromosome assembly (Chr1-Chr8, Fig. 4A). The remaining two contigs (Chr9a and Chr9b) reflect two telomere-to-centromere assemblies based on the telomeric repeat sequences identified at one end and the transposable elements clusters at the other consisting predominantly of long terminal repeat (LTR) retrotransposons and DNA transposons, which are typical signatures of centromeres in fungi (23), suggesting that Chr9a and Chr9b represent two arms of the same chromosome (Fig. 4A). Genes were distributed evenly along the chromosomes with an average of 247 genes per Mbp, except for regions of low gene distribution at the centromeres (Fig. 4A). Predicted gene function cataloging using the cluster of orthologous groups (COGs) database showed the presence of all major categories although the largest group of genes was classified as ‘Function unknown’ (Fig. 4B). The quality of the *A. oligospora* TWF154 genome sequence is amongst the most contiguous and complete of the sequenced ascomycetes. General features of the genome assemblies for the *A. oligospora* wild isolate TWF154 and ATCC24927 (24) are presented in Table 1. For example, the number of scaffolds and N50 in our annotation of TWF154 (10 scaffolds and N50=5,390,931) were improved in comparison with ATCC24927 (215 and 2,037,373, respectively). Using RNAseq data from five different culture conditions as guidance and the proteomes of representative fungi (see Supplementary Information), the program funannotate predicted 12107 gene models in our improved *A. oligospora* genome assembly; in comparison, 11479 genes had previously been found for ATCC24927 (Table 1). The proteome is 97.7% complete based on the BUSCO (25) assessment using the Ascomycota dataset, which is comparable to other published reference fungal genomes. Furthermore, we also assembled and annotated a complete mitochondrial genome (Fig. 4C). Notably, the size of mitochondrial genome of TWF154 (∼161 kb) is much larger than that of the human (∼17 kb), *N. crassa* (∼65 kb) or *S. cerevisiae* (∼79 kb) mitochondrial genomes. The expansion of the mitochondrial genome of TWF154 and other fungal mitochondrial genomes, namely in NTF species (26–34) can be attributed to the presence of a substantial number of genes encoding homing endonucleases (HEG) of the LAGLI-DADG and GIY-YIG families in intronic and intergenic regions (Fig. 4C). It will be interesting to evaluate whether the distribution of introns and HEGs varies in diverse *Arthrobotrys* strains and, more generally, the role of these genes during fungal-nematode interactions.

Table. 1. Genomic features the two assemblies of *A. oligospora* genomes

**Table. 1. Genomic features the two assemblies of *A. oligospora* genomes**
| General features | ATCC24927 | TWF154 |
| --- | --- | --- |
| Size of assembled genome (bp) | 40,072,829 | 39,619,008 |
| Number of scaffolds | 215 | 10 |
| N50 scaffold size (bp) | 2,037,373 | 5,390,931 |
| N90 scaffold size (bp) | 691,077 | 2,932,219 |
| GC content (%) | 44.45 | 43.96 |
| Number of genes | 11479 | 12107 |
| Number of tRNA genes | 145 | 102 |
| Busco completeness (%) | 97.4 | 97.7 |

**Figure. 4.**
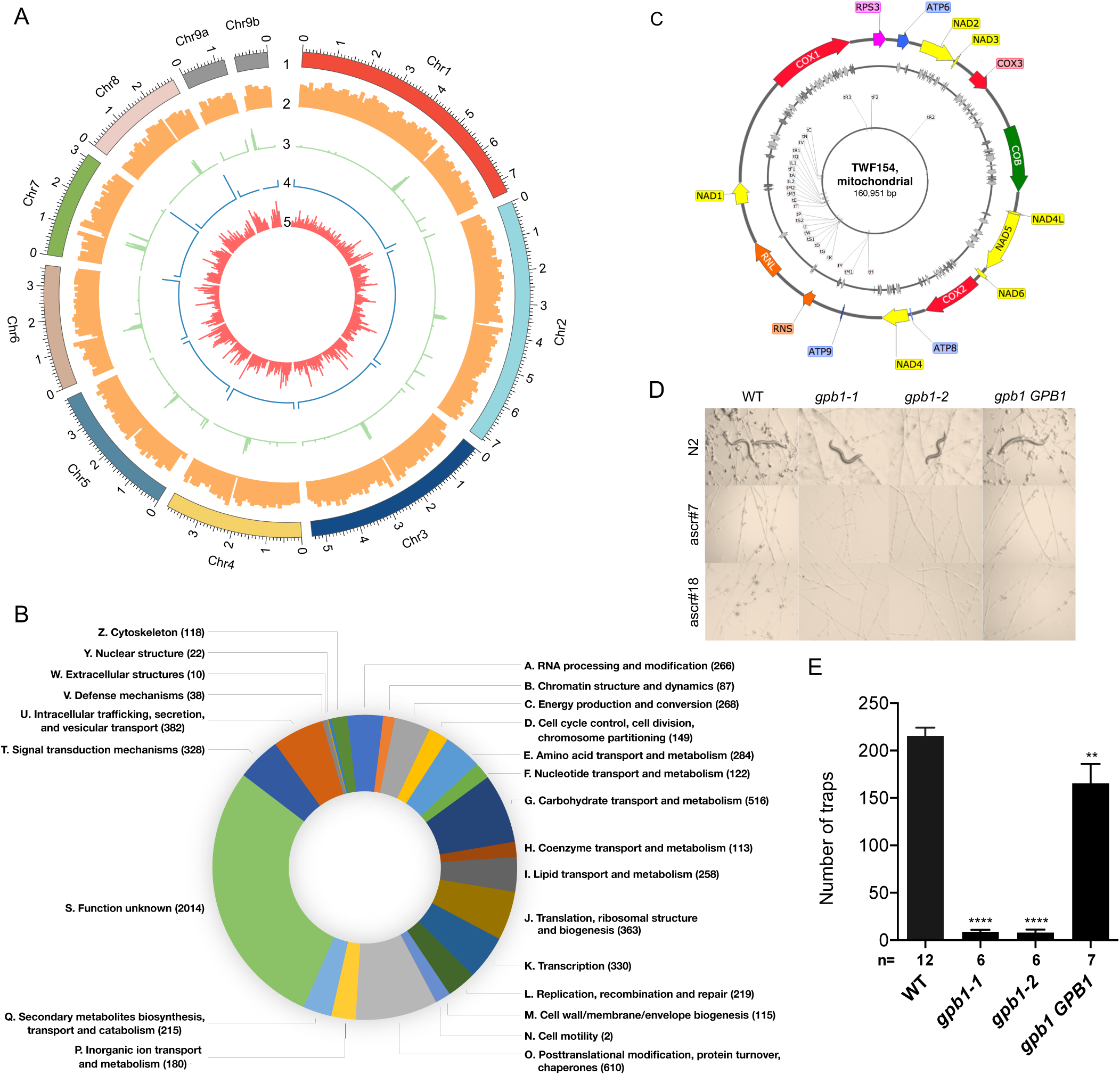
G protein signaling is required for prey sensing in *A. oligospora*. (A) Genome architecture of *A. oligospora* strain TWF154. Tracks (outer to inner) represent the distribution of genomic features: 1) positions (in Mb) of the ten TWF154 contigs, with numbers indicating the order of scaffolds size; 2) gene density (along a 100kb sliding window); 3) distribution of transposable elements (along 10kb sliding window); 4) telomere repeat frequency (along a1kb sliding window), showing that eight contigs have two telomeric ends; and 5), *A. oligospora* species-specific gene content (along a 100 kb sliding window). (B) Predicted function of genes in TWF154 genome cataloged using the cluster of orthologous groups (COGs) database. (C) Circular map of the mitochondrial genome of *A. oligospora* TWF154. Tracks (outer to inner) show: 1) annotation of mitochondrial DNA-encoded genes: subunits of NADH dehydrogenase/complex I (yellow), cytochrome *c* oxidase/complex IV (red) and ATP synthase/complex V (blue); apocytochrome *b COB* (green); ribosomal protein S3 *RPS3* (pink); ribosomal RNA genes *RNS* and *RNL* (orange); 2) approximate location of LAGLI-DADG (light gray) and GIY-YIG endonucleases (dark gray); 3) annotation of transfer RNA, where ‘t’ stands for ‘tRNA’ followed by the respective amino acid one letter code and number of copys. (D) Images of the traps induced by *C. elegans* lab strain N2 or ascarosides (ascr#7 and ascr#18) in the wild-type (TWF154), *gpb1* mutants or *gpb1 GPB1* rescued strains. Bar = 200 µm. (D) Quantification of the trap numbers induced by *C. elegans* wild-type strain N2 or (D) ascarosides (ascr#7 and ascr#18) in the wild-type (TWF154), *gpb1* mutants or *gpb1 GPB1* reconstituted strain (n shown along x-axis).

### G-protein signaling is required for prey-sensing in *A. oligospora*

We next performed functional studies of candidate genes that might be involved in sensing the nematodes. In *C. elegans*, ascarosides are sensed by G protein-coupled receptors (GPCR) (35, 36) and in fungi, G protein signaling and GPCRs are critical for sensing environmental stimuli, including sex pheromones, and are required for virulence and morphogenesis in various fungal pathogens (37, 38). We thus investigated if G protein signaling was involved in signaling in *A. oligospora* upon exposure to nematode cues. It had been suggested via chemical inhibitors that G proteins might be involved in the closure of the constriction ring of another nematode-trapping fungus *Drechslerella dactyloides*(39); however, the role of G protein signaling was never examined genetically. We thus identified a single G-protein *β* subunit gene (*GPB1*, EYR41_009480) and three G-protein alpha subunit genes (EYR41_010456, EYR41_011543, EYR41_011720) in the genome of *A. oligospora*. Since the G*α*s are known to have overlapped functions in many fungi(40), we decided to test the function of the G-protein *β* subunit first. We obtained two independent *gpb1* deletion mutants generated via homologous recombination and determined that the efficiency of homologous recombination was extremely low (∼3 %). The two *gpb1* mutants were strongly defective in trap morphogenesis in response to *C. elegans* and ascarosides (Fig. 4D, E). This phenotype was complemented by re-introducing the wild type copy of the *GPB1* gene (Fig. 4D, E). These results demonstrate that G protein signaling is required for ascaroside-sensing and/or trap formation in *A. oligospora* and that the ascaroside receptors in *A. oligospora* might be GPCRs.

### Conclusion

Our study revealed that NTF are widespread in soils of distinct ecological provenances, behave as generalist predators, are sympatric with diverse species of nematodes, and vary considerably in their spatial and temporal ability to capture the nematode prey. NTF hold great potential to be utilized as biological control agents in agricultural settings. However, only recently have the molecular mechanisms underlying the singular biology of these specialized predators began to be uncovered. For example, a recent work has generated a strain lacking the signaling scaffold protein Soft and shown that cell-cell fusion is required for ring closure in *Duddingtonia flagrans* (41). In *A. oligospora*, ATCC24927-background trapping-deficient mutants have been isolated: adhesin protein Mad1 (42), autophagy protein Atg8 (43), mitogen-activated protein kinase Slt2 (44), pH sensor PalH (45), NADPH oxidase NoxA (46), low-affinity calcium uptake system proteins Fig1 and Fig2 (47), Rab GTPase Rab7A (48), actin-associated protein Crn1 (49), Woronin body component Hex1 (50), malate synthase Mls (51), glycogen phosphorylase Gph1 (52), transcription factors VelB (53) and StuA (54), and microRNA processing protein Qde2 (55), to which we add the G-protein *β* subunit Gpb1. However, a mechanistic view of how these and other molecular players are interconnected during the interaction of *A. oligospora* with their prey is totally lacking. The establishment of a high-efficiency *A. oligospora* model strain (TWF154) and the generation of publicly available whole genome sequencing resources for an ample number of wild isolates will pave the way to the identification of novel loci underlying phenotypic traits and, more generally, to the fast advancement of research with NTF.

## Materials and Methods

### Isolation of nematodes and NTF from soil samples

Nematodes and NTF were isolated using the soil sprinkle method (56) with some modifications. Briefly, soil samples were sprinkled on low nutrient medium (LNM) and after 2-7 days, nematodes were transferred to nematode growth media (NGM) containing OP50 *Escherichia coli*. SSU rDNA sequences were PCR amplified directly from the nematodes using the single worm PCR protocol with primers G18S4a (5’-GCTCAAGTAAAAGATTAAGCCATGC-3’) and DF18S-8 (5’-GTTTACGGTCAGAACTASGGCGG-3’) (57, 58). NCBI BLAST (59) was used to determine the identity of the nematode isolates to species level (if the identity was higher than or equal to 97% when compared to a known species), or to the genus level (if the identity was lower than 97%). Once nematodes had been isolated from the LNM plates, *C. elegans* were added to the same plates to induce trap formation in the NTF. After 3-7 days, the plates were examined under a stereo microscope and single conidia of the NTF were isolated and transferred individually to potato dextrose agar (PDA) to establish pure cultures. ITS regions were directly amplified from fungal hyphae by PCR with the universal primers ITS1 5’-TCCGTAGGTGAACCTGCGG-3’ and ITS4 5’-TCCTCCGCTTATTGATATGC-3’ following the single worm PCR protocol (58, 60). Species identity was assigned if the ITS sequence identity was 97% or higher from NCBI BLAST (59) analyses. Strains of NTF and nematodes used in this study are listed in Table S4. *C. elegans* animals employed in this study were maintained following standard procedures (57).

### Phenotypical characterization

To quantify trap formation, fungal isolates were grown on LNM medium for 5 days, transferred to a fresh LNM plate and, after ∼48 hours (25°C, dark), exposed to 30 N2 *C. elegans* L4 stage larvae for 6 hours (after which the animals were removed with a pick) or 1 µl of 1 mM ascaroside #18 (61) that was added ∼2 mm away from the border of hyphal tips. Ascaroside #18 was synthesized as reported (62) and was of greater than 98% purity. Micrographs were taken after 24 hours after the addition of nematodes or ascaroside using a Nikon SMZ 745T stereo microscope. For each plate, 3 images were randomly taken within 0.5 cm from the edge of the plate using a 40x magnification, and the sum of traps in the 3 images was recorded. To estimate the survival rate of *C. elegans* after exposure to *A. oligospora*, the number of moving nematodes was computed every 2 hours for a total of 12 hours using a stereo microscope. Nematodes crawling on the wall of the plate, and therefore not in contact with the fungal hyphae, were excluded. Survival rates are presented as the percentage of moving worms over total number of worms. The time for trap emergence after contact with nematodes was measured in the same conditions as described for the previous assays but 5 cm LNM plates, 3 days of growth (25°C, dark) and 300 adult N2 *C. elegans* were used. A ZEISS Stemi 305 microscope with a ZEISS Axiocam ERc 5s camera was used to automatically capture images of the colony edge every 60 seconds for a total of 12 hours. Between 3-4 time-lapse videos were taken for each strain consisting of 2560×1920 pixels images representing an area of about 5.7×4.3 mm of the edge of the fungal colony. The time-lapse videos were visually inspected to record the time point at which the first trap arose in the imaged areas. For the competition assay, a Hygromycin-B (HygB)-resistant TWF154 strain was constructed by transforming a HygB resistance cassette (amplified from vector pAN7-1 (63) into TWF154 protoplasts (please see Supplementary Information). The HygB-resistant TWF154 strain showed no defect in growth, conidiation, and trap morphogenesis compared to wild type. Single spores of both HygB-resistant TWF154 and TWF106 (HygB sensisitive) were collected and transferred together to the center of replicated 5 cm LNM plates. After 4 days of growth, 300 adult N2 *C. elegans* were added to half of the plates and all fungal colonies were further grown for 3 more days to allow for trapping and digestion of the nematodes by the fungi. A spore suspension was then collected from the plates by adding 2 ml _dd_H2O and scratching the surface of the fungal colony, followed by centrifugation (1 min, 13200 rpm) and removal of the supernatant, yielding ∼0.2 ml highly concentrated spore solutions. Between 8-12 µl of each spore suspension were added to 9 cm PDA plates and PDA with 50 µl/ml HygB, respectively; all plates were treated with 90 µl 11 mg/ml ampicillin before the spore solution was added to prevent bacterial contamination. After 24 hours, the germinated spores in each plate were counted using a ZEISS Stemi 305 microscope. The competition rate was computed as the average number of germinated spores in the HygB plates over the average number of germinated spores in the PDA plates.

### TWF154 genome assembly and annotation, and phylogenetic analyses of nematode-trapping fungi

The TWF154 reference genome was sequenced and assembled from ∼0.5 million long reads with an average length of 11258 bp sequenced from the Pacbio RSII platform and polished with 18 million 250 bp Illumina reads. Annotation was done using funannotate (64) and the pipeline described in https://funannotate.readthedocs.io/en/latest/tutorials.html. Annotation of the mitochondrial genome was performed using a combination of MFannot (65) and MITOS2 (66). Details can be found in the supplementary information. The Neighbor-Joining algorithm of MEGA7 or MEGAX was used to construct maximum likelihood phylogenetic trees. The tree of 16 different NTF species isolated from soil samples was constructed based on ITS nucleotide sequences. *Tuber melanosporum* was the outgroup. The tree of the 20 *A. oligospora* isolates was constructed using 500 random single-copy orthologs from ATCC24927, TWF154 reference genome and 18 additional wild isolate genomes*. Dactylellina haptotyla* (strain CBS200) was used as the outgroup. The genomes of 18 wild isolates of *A. oligospora* were assembled from 18 million 250 bp Illumina reads using AAFTF (67) and annotated using funannotate (64). Further details about the assembly and annotation of the fungal isolates, as well as tree construction can be found in the supplementary information.

### Identification and deletion of the *GPB1* homolog in *A. oligospora*

The *GPB1* homolog of *A. oligospora* was identified with Blast2GO 5 Pro (68) by performing a Local BLAST using the amino acid sequence of Gpb1 orthologs of *N. crassa* (Uniprot ID: Q7RWT0), *Aspergillus nidulans* (Q5BH99), *Fusarium oxysporum* (Q96VA6), and *S. cerevisiae* (A6ZP55) as the query and the proteome of the T727 strain of *A. oligospora* as the database. *GPB1* was deleted by a homologous recombination-based strategy (69). Detailed methods including primers used for generating the *gpb1* deletion mutant are described in the supplementary information.

## Acknowledgements

The authors thank John Wang, Jun-Yi Leu and Sheng-Feng Shen for their helpful comments and suggestions about this work. We also than Ling-Mei Hsu and A-Mei Yang for their technical assistance. This work was supported by the start-up fund of Academia Sinica and Taiwan Ministry of Science and Technology 106-2311-B-001-039-MY3 to YPH.

## Supplemental Information

### Supplemental Materials and Methods

#### TWF154 genome Assembly and annotation

We *de novo* assembled the genome of strain TWF154 using ∼0.5 M Seqs long reads with an average length of 11258 bp, sequenced with the Pacbio RSII platform and accounting for a 131X coverage. First, preliminary assemblies using two different assembling pipelines were constructed. The first preliminary assembly was generated using Canu v.1.7 (1) with parameters: genomeSize=40m useGrid=false maxThreads=8. This assembly was then polished using Quiver and the original PacBio data. The polished Canu assembly was 39807769 nt divided in 16 contigs with a N50 of 3,921,408. In parallel to the Canu assembly we made a second assembly using the same PacBio data and a different assembler, Falcon (pb-assembly_0.0.2, (2). In order to run Falcon, we adapted the configuration file for fungal genomes assembling provided at https://pb-falcon.readthedocs.io/en/latest/parameters.html. We changed the Genome_size parameter to 40000000 and the memory and output options to the requirements of our local server. The assembly generated with Falcon was polished using Quiver, from the genomicconsensus v2.3.2 package of Bioconda (2), resulting in a final genome of 39,637,123nt distributed in 15 contigs and N50 of 3962661.

While the assembly obtained with Canu contained more complete telomeric regions, the Falcon assembly was more contiguous. Consequently, we decided to merged both assemblies using Quickmerge (3). This program allows to fill in the gaps of a reference assembly using the information contained in a donor assembly. In theory, the most trustworthy assembly should be the one used as the donor but in our case, we found strong points in both of our preliminary assemblies and therefore tried both combinations, *i.e.*, Canu as reference and Falcon as donor and *vice versa*. After merging the assemblies, two assemblies, one consisting of 15 contigs and a second of 13 contigs were obtained, both of them with higher N50 than both preliminary assemblies.

We searched for telomeric regions in the contigs of the merged assemblies by looking for TTAGGG repeats. The assembly where Canu was used as the reference and Falcon the donor, consisting of 13 contigs, was selected. This assembly contained in total 18 telomeric regions, 14 of which were found at the beginning and end of 7 of the contigs. Redundant contigs analysis with MUMMER4 (4) led us to remove 2 of the contigs since they were contained in a bigger contig. We merged two contigs that contained only one telomeric region and that presented more than 15kb (17633bp) of overlapping sequence. Consequently, we merged these two contigs. Finally, we polished the resulting assembly in two consecutive steps, first, by running Quiver on the final assembly and then using Illumina reads to further polish the assembly with Pilon (5). In total, about 18 M Seqs illumina reads of 251 bp were used for the pilon polishing, corresponding to a coverage of 112x coverage.

In order to annotate the assembled genome of TWF154 we used funannotate (6) and the pipeline described in https://funannotate.readthedocs.io/en/latest/tutorials.html. Consequently, we first softmasked the repetitive regions of TWF154 genome using funnanote mask. Following, we used funannotate train with RNA-seq data and masked genome to generate preliminary gene models. RNA was extracted from the same *A. oligospora* strain (28 MSeq of 76bp length). We then used funnanotate predict where the outputs from the training step and other gene prediction inputs are used to generate consensus gene models. These gene models were further corrected using the RNA-seq data by using the command funannotate update, which also adds UTR data to the predictions. In order to perform additional functional annotation for our fixed gene models, we ran in parallel and using the funannotate remote option: InterProScan, Eggnog-mapper and antiSMASH yielding information on InterPro terms, GO ontology and fungal transcription factors, Eggnog annotations and COGs, and secondary metabolites, respectively. The latter outputs were used as inputs for funannotate which additionally identifies Pfam domains, CAZYmes, secreted proteins, proteases (MEROPS), and BUSCO groups. The results of the funnanotate annotation consist of several files combining the annotation data obtained from all the above mentioned steps, including genbank format and others required for NCBI submission.

The final assembled and annotated genome of *A. oligospora* TWF154 was uploaded to NCBI and is available with accession number: SOZJ00000000.

##### Annotation of the TWF154 mitochondrial genome

A circular DNA contig was obtained from the Canu assembly step of whole genome sequencing analysis. We observed a putative large duplication by aligning this sequence with itself and suspected this to be an artifact due to the circular nature of the mitochondrial DNA and overlap between assembled PacBio reads. Our hypothesis was confirmed by aligning the contig with various fungal mitochondrial genomes and the large duplicated region was manually removed to obtain the final mitochondrial genome of TWF154, which had an identical size to the mitochondrial genome of ATCC24927 (7). Annotation of the mitochondrial genome was performed using a combination of MFannot (8) and MITOS2 (9). The annotated mitochondrial genome was uploaded to NCBI (submission ID 2280642).

#### *A. oligospora* wild isolates genome assembly and annotation

The genome of 18 wild isolates of A. oligospora was assembled using Automatic Assembly For The Fungi (AAFTF) (10). AAFTF was first used to trim 18 million 250 bp Illumina reads. The filtered reads were then used to make preliminary assemblies using the Spades option of AAFTF assemble. Contaminants and segments of vector origin were removed from the assemblies using the soursmash and vecscreen functions of AFFTF, respectively. Duplicated scaffolds were removed using the AAFTF rundup and the resulting assemblies were finally polished using the PILON function of AAFTF.

The final genomes were annotated as described in the previous section and using the same RNA-seq data for training as TWF154. The genome statistics of the wild isolate assemblies can be found in Supplementary Table 1 together with their NCBI accession numbers.

**Supplementary Table 1.** Genome summary of *A. oligospora* wild isolates and accession numbers.

| Wild isolate | Assembly Size | Scaffolds | N50 | Genes | Accession number |
| --- | --- | --- | --- | --- | --- |
| TWF102 | 40,306,818 bp | 425 | 272,178 bp | 12,028 | WIQW00000000 |
| TWF103 | 39,840,834 bp | 555 | 253,865 bp | 12,021 | WIQX00000000 |
| TWF106 | 40,488,918 bp | 661 | 197,328 bp | 11,725 | WIWS00000000 |
| TWF128 | 39,491,435 bp | 296 | 303,590 bp | 12,091 | Processing (SUB6434522) |
| TWF132 | 39,728,578 bp | 301 | 363,783 bp | 12,104 | Processing (SUB6433031) |
| TWF173 | 39,919,627 bp | 518 | 368,284 bp | 11,560 | Processing (SUB6438852) |
| TWF191 | 40,589,086 bp | 451 | 264,791 bp | 11,418 | WIPF00000000 |
| TWF192 | 40,487,552 bp | 453 | 334,966 bp | 11,542 | WIPG00000000 |
| TWF217 | 39,480,237 bp | 273 | 356,212 bp | 12,080 | Processing (SUB6437927) |
| TWF225 | 39,453,384 bp | 305 | 413,125 bp | 12,096 | WIWR00000000 |
| TWF569 | 39,905,426 bp | 482 | 275,809 bp | 11,982 | WIRA00000000 |
| TWF594 | 39,943,739 bp | 441 | 303,780 bp | 11,885 | WIRB00000000 |
| TWF679 | 40,193,099 bp | 539 | 236,485 bp | 11,482 | WIWT00000000 |
| TWF703 | 40,031,381 bp | 525 | 251,714 bp | 11,306 | WIQZ00000000 |
| TWF706 | 40,300,755 bp | 472 | 305,738 bp | 12,015 | WIQY00000000 |
| TWF751 | 39,943,019 bp | 335 | 280,041 bp | 12,123 | WIWQ00000000 |
| TWF788 | 40,740,697 bp | 364 | 224,796 bp | 11,646 | Processing (SUB6439089) |
| TWF970 | 39,580,990 bp | 193 | 669,589 bp | 11,625 | Processing (SUB6438993) |

#### Phylogenetic tree of *A. oligospora* wild isolates

The annotated genomes of the 18 wild isolates, TWF154, ATCC24927 and the outgroup *Dactylellina haptotyla* (strain CBS200) were used to construct the phylogenetic tree as follows. First, the compare function of funannotate was used to find orthologs among the different genomes by calling ProteinOrtho5 (11) with selfblast and blastp options. Funannotate selects from all the orthologs only single copy orthologs with hits in the BUSCO database and clusters them in orthology groups. 500 randomly selected orthologs are then aligned using mafft (12) and passed to trimAl (13) to remove poorly aligned regions. We used the trimmed alignment file as input for the Neighbor-Joining algorithm of MEGAX (Molecular Evolutionary Genetics Analysis version 10.1.1).

#### *A. oligospora GPB1* deletion and rescue

A homologous recombination method was employed to delete the *A. oligospora GPB1* gene (EYR41_ 009480). An overlap PCR-based construct (donor DNA) was obtained by fusing the 2 kb long sequences flanking the open reading frame of *GPB1* (amplified from genomic DNA of the T727) to a hygromycin-B resistance cassette (amplified from vector pAN7-1 (14)). The construct was introduced into protoplasts of the T727 strain via PEG-mediated transformation. Generation of protoplasts in *A. oligospora* was carried out based on previously described protocols with slight modifications (15, 16). Approximately 5 x 10^6^ of conidia were cultured in 100 ml of liquid PDB for 24 hours at 25°C, 200 rpm. Mycelia were collected by centrifugation, washed with MN buffer (0.3 M MgSO_4_, 0.3 M NaCl), mixed with 5 ml of 100 mg/ml VinoTastePro lytic enzyme mix in MN buffer and incubated for 6 to 8 hours at 30°C, 200 rpm. Protoplasts were separated from mycelial debris by filtering through two layers of miracloth tissue and washed with STC buffer (1.2 M sorbitol, 50 mM CaCl_2_, 10mM Tris-HCl pH 7.5). The protoplasts were then resuspended in STC buffer to a final concentration of 1 x 10^7^ protoplasts/ml and used immediately for PEG-mediated transformation. For transformations, ∼10^6^ protoplasts were gently mixed with 5 µg of donor DNA and incubated on ice for 30 mins, after which five volumes of PTC (40% w/v PEG 3350, 10 mM Tris-HCl pH 7.5, 50 mM CaCl_2_) were added and incubated at room temperature for 20 mins. Then, 50 ml of molten PDA agar at 45°C containing 100 µg/ml Hygromycin B was added to the protoplast mixture and poured into Petri dishes. After 7 days at 25°C, transformants grown on selective medium were screened for gene replacement by rapid genomic extraction and PCR. Briefly, a small cluster of aerial hyphae was added to PCR tubes containing 20 µl of DNA lysis buffer (50mM KCl, 10 mM Tris pH8.3, 2.5 mM MgCl_2_, 0.45% Nonidet P-40, 0.045% Tween-20, 0.01% (w/v) gelatinin) that were placed in a thermal cycler and treated with the following program for gDNA extraction: 65°C (45 mins), 95°C (7 mins), 10°C (10 mins). Two µl were then used as template for PCR reactions. For the construction of a *GPB1* rescue strain, a wild type copy of *GPB1* was amplified from T727 genomic DNA (2 kb upstream and 1 kb downstream of the open reading frame) and cloned into pRS41N containing a nourseothricin-resistance cassette. The resulting vector was transformed into the *gpb1* mutants. All the primers used for constructing gene deletion and complementation are listed in Supplementary Table 5.

#### Statistical analysis

GraphPad Prism was used to draw plots and perform statistical analysis (tests are indicated in the respective figure legends).

**Supplementary Table 2.** Wild isolates of nematodes and nematode-trapping fungi in this study.

| Taiwan |  |  |  |
| --- | --- | --- | --- |
| City | Sampling Site | Worm Species | NTF Species |
| Chiayi | Alishan 1 | *Acrobelloides nanus*, *Bunonema reticulatum*, *Teratocephalus terrestris*, *Acrobelloides thornei*, *Choriorhabditis cristata*, | *Arthrobotrys oligospora* |
| Hualien | Pingfengshan 1 | *Pristionchus* sp., *Rhabditophanes* sp., *Acrobelloides apiculatus* | N/A |
|  | Pingfengshan 2 | *Oscheius tipulae* | N/A |
|  | Pingfengshan 3 | *Mesorhabditis* sp. | N/A |
|  | Pingfengshan 4 | *Acrobelloides apiculatus*, *Pristionchus* sp. | N/A |
|  | Pingfengshan 5 | *Acrobelloides thornei*, *Acrobelloides apiculatus* | N/A |
|  | Pingfengshan 6 | *Acrobelloides* cf. | N/A |
|  | NDHU 1 | N/A | *Arthrobotrys oligospora* |
|  | Liyu Lake 1 | *Mesorhabditis* sp. | *Arthrobotrys thaumasia* |
|  | Butterfly Valley 1 | *Oscheius tipulae*, *Mesorhabditis* sp. | *Arthrobotrys thaumasia*,<br>*Arthrobotrys vermicola* |
| Ilan | Fushan 1 | *Oscheius tipulae*, *Pelodera* sp. | *Arthrobotrys oligospora*,<br>*Arthrobotrys megalosporum*,<br>*Arthrobotrys thaumasia* |
|  | Ming Chi 1 | *Oscheius tipulae*, *Rhabditis blumi* | *Monacrosporium geophyropaga* |
|  | Taipingshan 1 | *Oscheius tipulae* | N/A |
|  | Taipingshan 2 | *Oscheius tipulae* | N/A |
|  | Toucheng Farm 1 | *Oscheius tipulae*, *Oscheius chongmingensis* | *Arthrobotrys oligospora*,<br>*Arthrobotrys musiformis*,<br>*Arthrobotrys thaumasia* |
| Miaoli | Zhunan 1 | *Caenorhabditis sinica* | *Arthrobotrys oligospora* |
| Nantou | Hehuanshan 1 | *Panagrolaimus detritophagus*, *Acrobeloides apiculatus* | N/A |
|  | Huisun 1 | *Oscheius tipulae*, *Aphelenchoides* sp.,<br>*Acrobeloides* sp. | *Arthrobotrys musiformis*,<br>*Arthrobotrys megalosporum*,<br>*Arthrobotrys* sp. |
|  | Huisun 2 | *Oscheius tipulae* | *Arthrobotrys* sp., *Arthrobotrys megalosporum* |
|  | Huisun 3 | *Oscheius tipulae*, *Pristionchus pacificus* | *Arthrobotrys thaumasia* |
|  | Huisun 4 | *Oscheius tipulae*, *Rhabditis myriophila* | *Arthrobotrys megalosporum*,<br>*Arthrobotrys musiformis*,<br>*Arthrobotrys oligospora* |
|  | Huisun 5 | *Acrostichus* sp. | *Arthrobotrys* sp. |
|  | Sitou 1 | *Rhabditis myriophila* | *Arthrobotrys sinensis* |
|  | Mingjian 1 | N/A | *Arthrobotrys* sp., *Arthrobotrys oligospora* |
|  | Highland Experimental Farm 1 | *Acrobeloides apiculatus*, *Ceratoplectus* cf.,<br>*Chiloplacus* sp., *Oscheius tipulae*,<br>*Diplogasteroides* sp., *Plectus andrassyi*,<br>*Aporcelaimellus* sp., *Rhabditella axei*, *Pristionchus hoplostomus*, *Demaniella* sp., *Pelodera teres*,<br>*Caenorhabditis elegans*, *Pelodera* sp.,<br>*Mesorhabditis spiculigera*, *Oscheius* sp.,<br>*Pristionchus* sp., *Aphelenchoides* sp.,<br>*Caenorhabditis* sp., *Sheraphelenchus susus* | *Arthrobotrys oligospora*,<br>*Arthrobotrys sinensis*,<br>*Arthrobotrys cookedickinsonianus* |
|  | Highland Experimental Farm 2 | *Oscheius tipulae*, *Plectus andrassyi*, *Alaimus* sp.,<br>*Aphelenchoides* sp., *Acrobeloides apiculatus*,<br>*Mesorhabditis* sp., *Diplogastrellus* sp.,<br>*Aporcelaimellus* sp., *Cervidellus alutus*,<br>*Ceratoplectus* cf. | *Arthrobotrys musiformis* |
|  | Highland Experimental Farm 3 | *Oscheius tipulae*, *Acrobeloides* sp., *Acrobeloides apiculatus*,<br>*Mesorhabditis* sp., *Aphelenchoides* sp.,<br>*Aphelenchoides bicaudatus*, *Plectus andrassyi*,<br>*Wilsonema otophorum* | *Arthrobotrys oligospora*,<br>*Arthrobotrys thaumasia* |
| New Taipei | Er Zi Ping 1 | *Oscheius tipulae*, *Mesorhabditis* sp. | *Arthrobotrys* sp., *Arthrobotrys musiformis* |
|  | Er Zi Ping 2 | *Oscheius tipulae*, *Bursaphelenchus* sp.,<br>*Caenorhabditis briggsae*, *Oscheius myriophila*,<br>*Bursaphelenchus fungivorus* | *Arthrobotrys megalosporum*,<br>*Arthrobotrys javanica* |
|  | Great Roots 1 | Unidentified nematode species | N/A |
|  | Wugu 1 | *Rhabditis rainai* | N/A |
|  | Qingtiangang 1 | *Oscheius tipulae*, *Diploscapter coronatus*, *Acrobeloides* cf., *Cervidellus vexilliger* | *Arthrobotrys thaumasia* |
|  | Qingtiangang 2 | *Acrobeloides apiculatus* | *Arthrobotrys thaumasia* |
|  | Songluo Lake 1 | *Acrobeloides nanus*, *Caenorhabditis* sp., *Acrobeloides bodenheimeri* | *Arthrobotrys thaumasia* |
|  | Wulai 1 | *Oscheius tipulae*, *Caenorhabditis wallacei*, *Protorhabditis* sp., *Caenorhabditis briggsae*, *Caenorhabditis tropicalis*, *Acrostichus* sp. | *Arthrobotrys vermicola*,<br>*Arthrobotrys oligospora*,<br>*Arthrobotrys javanica* |
|  | Xindian 1 | Unidentified nematode species | N/A |
|  | Xiaoyoukeng 1 | N/A | N/A |
|  | Mt. Chahu 1 | *Acrobeloides apiculatus*, *Cervidellus alutus*, *Aphelenchus avenae*, *Cervidellus alutus* | *Arthrobotrys microscaphoides*,<br>*Arthrobotrys thaumasia* |
| Penghu | Bird Island 1 | *Mesorhabditis* sp. | *Drechslerella anchonia* |
|  | Penghu 1 | *Diploscapter coronatus*, *Rhabditis rainai*, *Mesorhabditis* sp., *Acrobeloides nanus* | *Arthrobotrys oligospora*,<br>*Arthrobotrys* sp. |
|  | Penghu 2 | N/A | N/A |
|  | Basalt 1 | *Acrobeloides nanus* | *Arthrobotrys* sp. |
| Taichung | Daxueshan 1 | *Acrobeloides apiculatus*, *Mesorhabditis* sp. | *Drechslerella coelobrocha*,<br>*Monacrosporium cionopagum* |
|  | Daxueshan 2 | *Oscheius* sp., *Pristionchus* sp., *Pristionchus fissidentatus*, *Alaimus* sp., *Choriorhabditis cristata*, *Wilsonema schuurmansstekhoveni*, *Plectus minimus*, *Acrobeloides nanus*, *Aporcelaimellus obtusicaudatus* | N/A |
|  | Daxueshan 3 | *Acrobeloides nanus*, *Oscheius* sp., *Wilsonema schuurmansstekhoveni*, *Pristionchus* sp., *Alaimus* sp., *Choriorhabditis cristata*, *Bunonema franzi*, *Teratocephalus terrestris* | *Arthrobotrys musiformis* |
|  | Lingmingshan 1 | *Acrobeloides apiculatus* | N/A |
|  | Lingmingshan 2 | *Acrobeloides apiculatus* | N/A |
|  | Lingmingshan 3 | *Acrobeloides apiculatus* | N/A |
|  | Lingmingshan 4 | Unidentified nematode species | N/A |
|  | Wufeng 1 | *Oscheius tipulae*, *Pseudacrobeles* sp., *Acrobeloides* cf., *Acrobeloides apiculatus*, *Mesorhabditis spiculigera* | *Arthrobotrys thaumasia* |
|  | Wufeng 2 | *Mesorhabditis* sp., *Aporcelaimellus* cf., *Acrobeloides apiculatus*, *Panagrolaimus trilabiatus* | *Arthrobotrys thaumasia* |
|  | Wufeng 3 | *Ecumenicus monohystera*, *Acrobeloides apiculatus*, *Panagrolaimus* sp., *Mesorhabditis* sp., *Eumonhystera filiformis* | *Arthrobotrys thaumasia* |
|  | Wuling Farm 1 | *Oscheius tipulae* | *Arthrobotrys oligospora*, *Monacrosporium haptotylum* |
|  | Wuling Farm 2 | *Oscheius tipulae*, *Acrobeloides apiculatus*, *Rhabditis* sp., *Pelodera teres* | *Arthrobotrys amerospora* |
|  | Wuling Farm 3 | *Acrobeloides apiculatus* | *Arthrobotrys thaumasium* |
|  | Wuling Farm 4 | *Oscheius tipulae* | *Arthrobotrys musiformis*, *Arthrobotrys oligospora* |
|  | Wuling Farm 5 | Unidentified nematode species | *Arthrobotrys oligospora* |
| Taipei | Academia Sinica 1 | *Mesorhabditis* sp., *Panagrolaimus subelongatus*, *Oscheius tipulae*, *Cervidellus vexilliger*, *Oscheius myriophilus* | *Arthrobotrys thaumasia* |
|  | Academia Sinica 2 | *Rhabditis colombiana*, *Rhabditis rainai* | *Arthrobotrys thaumasia*, *Arthrobotrys oligospora*, *Arthrobotrys musiformis* |
|  | Nangang 1 | *Oscheius tipulae*, *Rhabditis colombiana*, *Oscheius carolinensis* | *Arthrobotrys oligospora*, *Arthrobotrys musiformis* |
|  | Neigou 1 | *Pelodera cylindrica*, *Rhabditis myriophila*, *Rhabditis rainal*, *Caenorhabditis sinica*, *Rhabditinae* sp. | *Arthrobotrys oligospora*, *Arthrobotrys musiformis* |
|  | Taipei 1 | N/A | *Arthrobotrys javanica*, *Arthrobotrys thaumasia* |
| Taoyuan | Lalashan 1 | *Panagrellus redivivus*, *Caenorhabditis* sp., *Sheraphelenchus sucus*, *Caenorhabditis briggsae*, *Cephaloboides* sp., *Oscheius tipulae* | *Arthrobotrys musiformis*, *Arthrobotrys oligospora* |
|  | Lalashan 2 | *Ceratoplectus* cf., *Protorhabditis* sp. | N/A |
|  | Lalashan 3 | *Diploscapter coronatus*, *Caenorhabditis briggsae*, *Panagrellus* sp., *Caenorhabditis* sp., *Sheraphelenchus sucus* | N/A |
| Taitung | Lanyu 1 | *Mesorhabditis* sp., *Aphelenchus avenae*, *Acrobeloides* sp., *Acrobeloides thornei*, *Cephalobus cubaensis*, *Cervidellus alutus* | *Arthrobotrys* sp. |
|  | Lanyu 2 | *Oscheius tipulae*, *Aphelenchus avenae*, *Cervidellus alutus*, *Mylonchulus hawaiiensis*, *Acrobeloides apiculatus* | *Arthrobotrys oligospora*, *Arthrobotrys guizhouensis*, *Arthrobotrys* sp., *Arthrobotrys vermicola*, *Arthrobotrys thaumasia* |
|  | Lanyu 3 | *Oscheius tipulae*, *Aphelenchus avenae*, *Acrobeloides apiculatus*, *Mesorhabditis* sp., *Cephalobus cubaensis*, *Panagrolaimus* sp., *Zeldia punctata*, *Acrobeloides thornei* | *Arthrobotrys* sp., *Arthrobotrys oligospora* |
| Tainan | Anding 1 | *Diploscapter coronatus*, *Acrobeloides apiculatus*, *Cephalobus cubaensis*, *Aphelenchoides fujianensis* | N/A |

**Supplementary Table 3.**
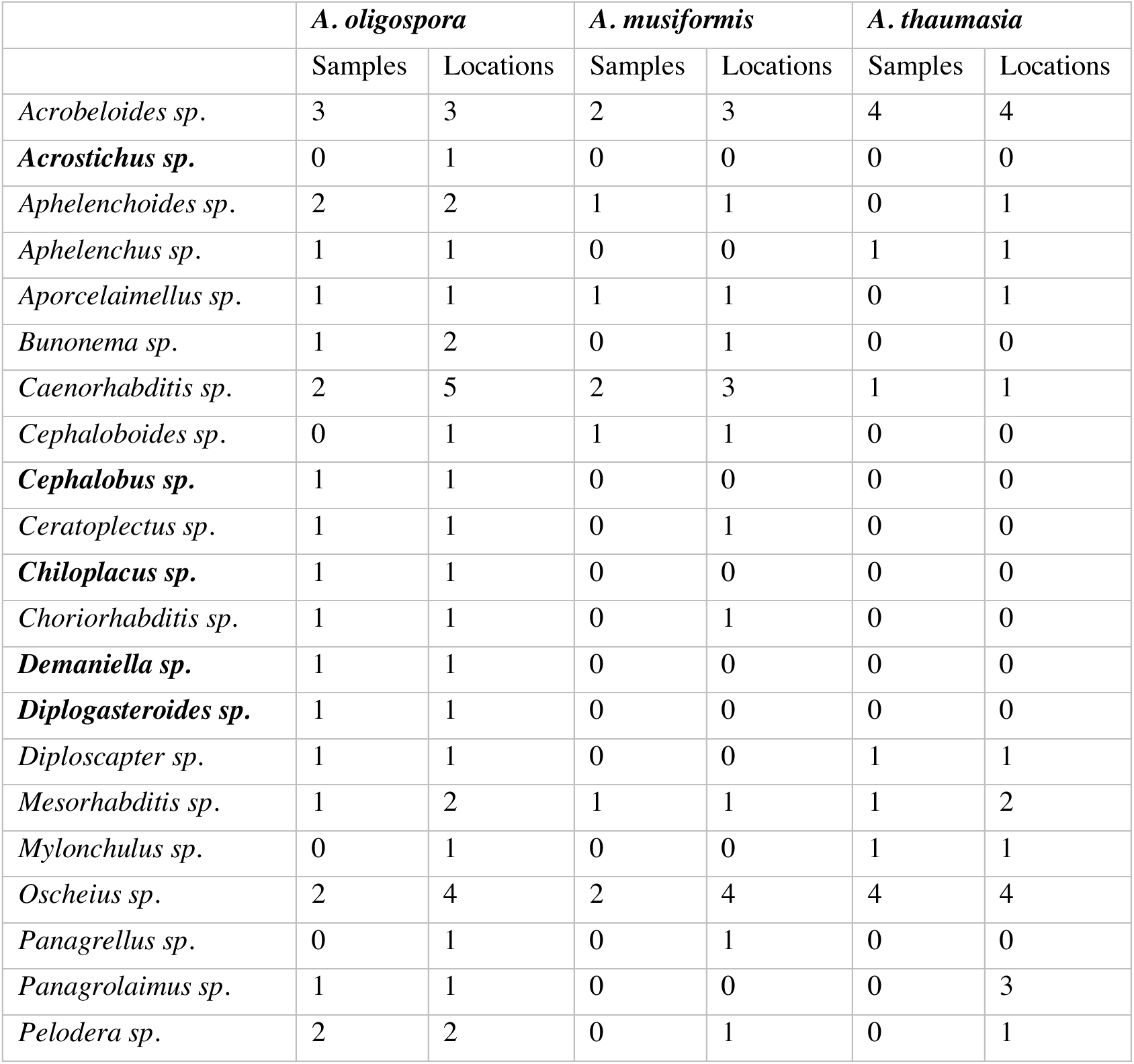
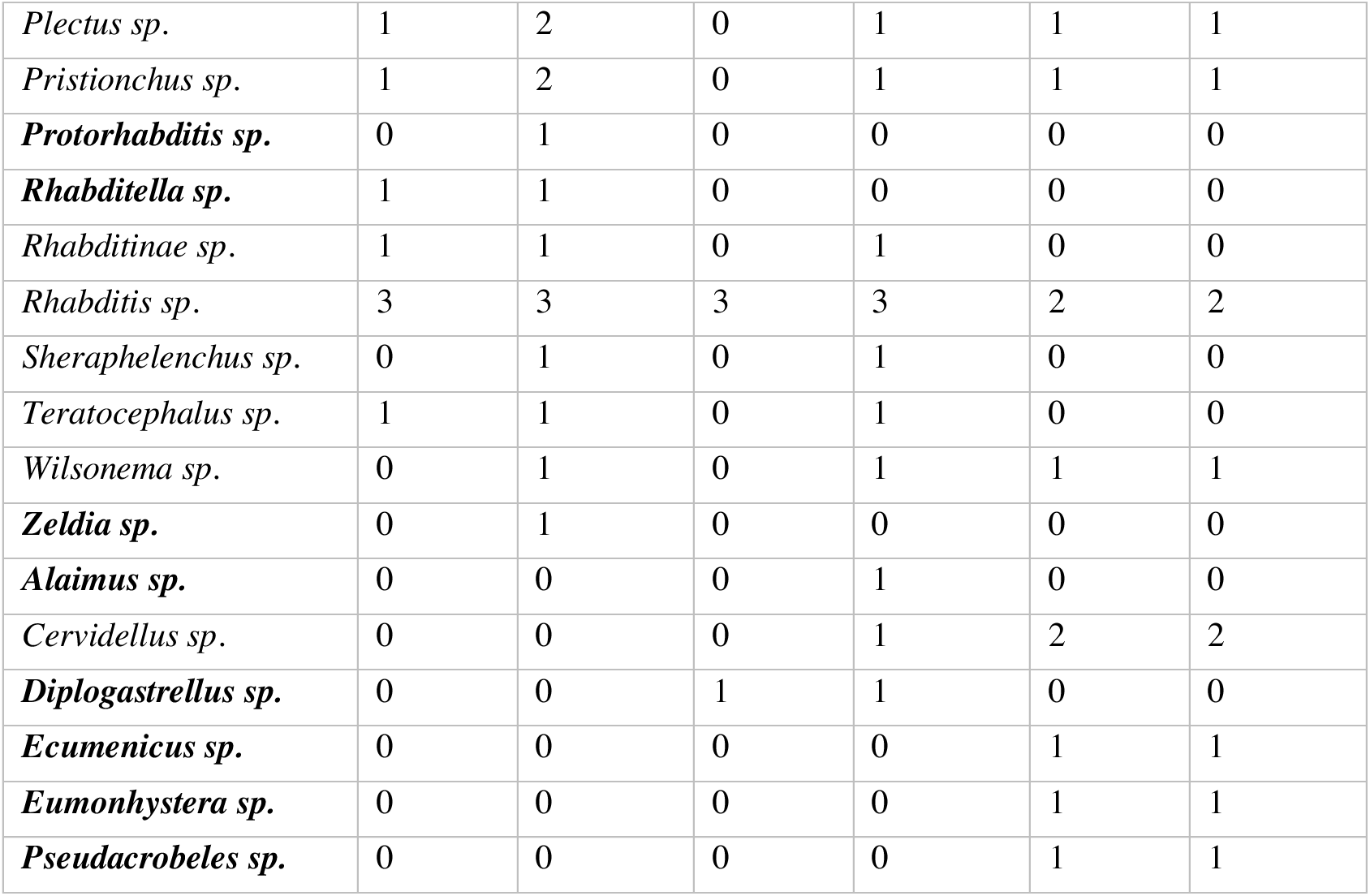
Species of nematodes sympatric with the three most common NTF identified in Taiwanese soil samples: *A. oligospora, A. musiformis* and *A. thaumasia*. The table shows, for each nematode and NTF species, the number of soil samples and locations where both organisms were observed coexisting. The species shown in bold represent nematode species coexisting with only one of the studied NTF species.

|  | *A. oligospora* |  | *A. musiformis* |  | *A. thaumasia* |  |
| --- | --- | --- | --- | --- | --- | --- |
|  | Samples | Locations | Samples | Locations | Samples | Locations |
| *Acrobeloides* sp. | 3 | 3 | 2 | 3 | 4 | 4 |
| ***Acrostichus* sp.** | 0 | 1 | 0 | 0 | 0 | 0 |
| *Aphelenchoides* sp. | 2 | 2 | 1 | 1 | 0 | 1 |
| *Aphelenchus* sp. | 1 | 1 | 0 | 0 | 1 | 1 |
| *Aporcelaimellus* sp. | 1 | 1 | 1 | 1 | 0 | 1 |
| *Bunonema* sp. | 1 | 2 | 0 | 1 | 0 | 0 |
| *Caenorhabditis* sp. | 2 | 5 | 2 | 3 | 1 | 1 |
| *Cephaloboides* sp. | 0 | 1 | 1 | 1 | 0 | 0 |
| ***Cephalobus* sp.** | 1 | 1 | 0 | 0 | 0 | 0 |
| *Ceratoplectus* sp. | 1 | 1 | 0 | 1 | 0 | 0 |
| ***Chiloplacus* sp.** | 1 | 1 | 0 | 0 | 0 | 0 |
| *Choriorhabditis* sp. | 1 | 1 | 0 | 1 | 0 | 0 |
| ***Demaniella* sp.** | 1 | 1 | 0 | 0 | 0 | 0 |
| ***Diplogasteroides* sp.** | 1 | 1 | 0 | 0 | 0 | 0 |
| *Diploscapter* sp. | 1 | 1 | 0 | 0 | 1 | 1 |
| *Mesorhabditis* sp. | 1 | 2 | 1 | 1 | 1 | 2 |
| *Mylonchulus* sp. | 0 | 1 | 0 | 0 | 1 | 1 |
| *Oscheius* sp. | 2 | 4 | 2 | 4 | 4 | 4 |
| *Panagrellus* sp. | 0 | 1 | 0 | 1 | 0 | 0 |
| *Panagrolaimus* sp. | 1 | 1 | 0 | 0 | 0 | 3 |
| *Pelodera* sp. | 2 | 2 | 0 | 1 | 0 | 1 |
| *Plectus sp.* | 1 | 2 | 0 | 1 | 1 | 1 |
| *Pristionchus sp.* | 1 | 2 | 0 | 1 | 1 | 1 |
| ***Protorhabditis sp.*** | 0 | 1 | 0 | 0 | 0 | 0 |
| ***Rhabditella sp.*** | 1 | 1 | 0 | 0 | 0 | 0 |
| *Rhabditinae sp.* | 1 | 1 | 0 | 1 | 0 | 0 |
| *Rhabditis sp.* | 3 | 3 | 3 | 3 | 2 | 2 |
| *Sheraphelenchus sp.* | 0 | 1 | 0 | 1 | 0 | 0 |
| *Teratocephalus sp.* | 1 | 1 | 0 | 1 | 0 | 0 |
| *Wilsonema sp.* | 0 | 1 | 0 | 1 | 1 | 1 |
| ***Zeldia sp.*** | 0 | 1 | 0 | 0 | 0 | 0 |
| ***Alaimus sp.*** | 0 | 0 | 0 | 1 | 0 | 0 |
| *Cervidellus sp.* | 0 | 0 | 0 | 1 | 2 | 2 |
| ***Diplogastrellus sp.*** | 0 | 0 | 1 | 1 | 0 | 0 |
| ***Ecumenicus sp.*** | 0 | 0 | 0 | 0 | 1 | 1 |
| ***Eumonhystera sp.*** | 0 | 0 | 0 | 0 | 1 | 1 |
| ***Pseudacrobeles sp.*** | 0 | 0 | 0 | 0 | 1 | 1 |

**Supplementary Table 4.** Wild isolates of nematode and NTF species used in this study.

| **Nematode Species** | **Source** | **NO.** |
| --- | --- | --- |
| *Caenorhabditis elegans* | *Caenorhabditis* genetics center | N2 |
| *Caenorhabditis tropicalis* | This study | TWN2033 |
| *Caenorhabditis wallacei* | This study | TWN2103 |
| *Caenorhabditis briggsae* | This study | TWN1756 |
| *Caenorhabditis sinica* | This study | TWN289 |
| *Oscheius tipulae* | This study | TWN1327, TWN1409,<br>TWN208, TWN272 |
| *Oscheius chongmingensis* | This study | TWN480, TWN1407 |
| *Oscheius dolichura* | This study | TWN1542 |
| *Mesorhabditis sp.* | This study | TWN1349 |
| *Rhabditis myriophila* | This study | TWN296 |
| *Rhabditis colombiana* | This study | TWN1422 |
| *Rhabditis rainai* | This study | TWN301 |
| *Pristionchus pacificus* | This study | TWN1985 |
| *Acrobeloides nanus* | This study | PHw8, Son1W1 |
| *Acrobelloides apiculatus* | This study | TWN1485 |
| *Panagrolaimus* sp. | This study | TWN1482 |
| *Cervidellus vexilliger* | This study | TWN1125 |
| **NTF Species** | **Source** | **NO.** |
| *Arthrobotrys oligospora* | This study | TWF95, TWF102, TWF103, TWF106, TWF107, TWF128, TWF132, TWF145, TWF149, TWF152, TWF154, TWF162, TWF173, TWF197, TWF217, TWF225, TWF537, TWF569, TWF679, TWF751, TWF788, TWF978, TWF706, TWF770, TWF996, TWF191, TWF192, TWF594, TWF703, TWF899, TWF915, TWF922, TWF925, TWF928, TWF970 |
| *Arthrobotrys thaumasia* | This study | TWF156, TWF258, TWF678, TWF547, TWF566, TWF586, TWF588, TWF735, TWF291, TWF1005, TWF233, TWF806 |
| *Arthrobotrys musiformis* | This study | TWF105, TWF140, TWF166, TWF556, TWF481, TWF571, TWF697, TWF791, TWF906, TWF784, TWF907 |
| *Arthrobotrys oligospora* | American Type Culture Collection | ATCC24927 |
| *Arthrobotrys vermicola* | This study | TWF730 |
| *Arthrobotrys guizhouensis* | This study | TWF891 |
| *Arthrobotrys amerospora* | This study | TWF531 |
| *Arthrobotrys javanica* | This study | TWF718 |
| *Arthrobotrys cookedickinsonianus* | This study | TWF920 |
| *Arthrobotrys sinensis* | This study | TWF918 |
| *Arthrobotrys megalosporum* | This study | TWF224 |
| *Arthrobotrys microscaphoides* | This study | TWF903 |
| *Monacrosporium haptotylum* | This study | TWF543 |
| *Monacrosporium gephyropaga* | This study | TWF768 |
| *Monacrosporium cionopagum* | This study | TWF492 |
| *Drechlerella coelobrocha* | This study | TWF491 |
| *Drechlerella anchonia* | This study | TWF766 |
| *Arthrobotrys oligospora*: *gpb1::hygR* | This study | TWF932 |
| *Arthrobotrys oligospora*: *gpb1::hygR* | This study | TWF933 |
| *Arthrobotrys oligospora*: *gpb1::hygR*;<br>*GPB1* | This study | TWF1140 |

**Supplementary Table 5.**
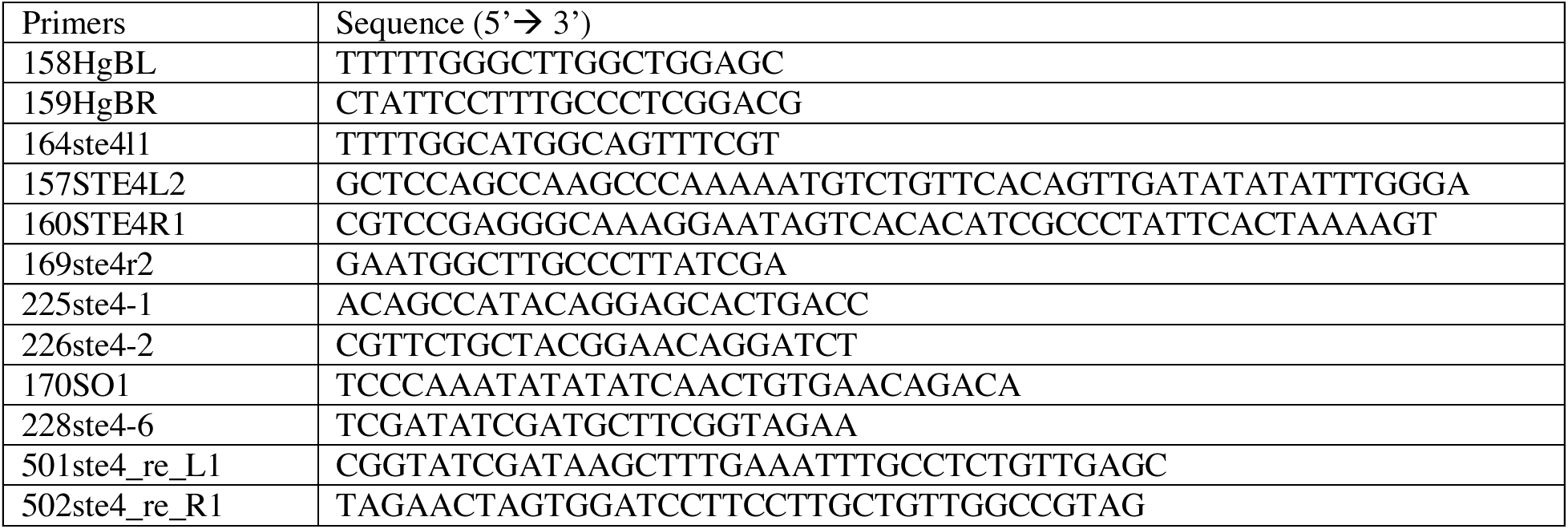
Primers used in this study.

**Figure S1.**
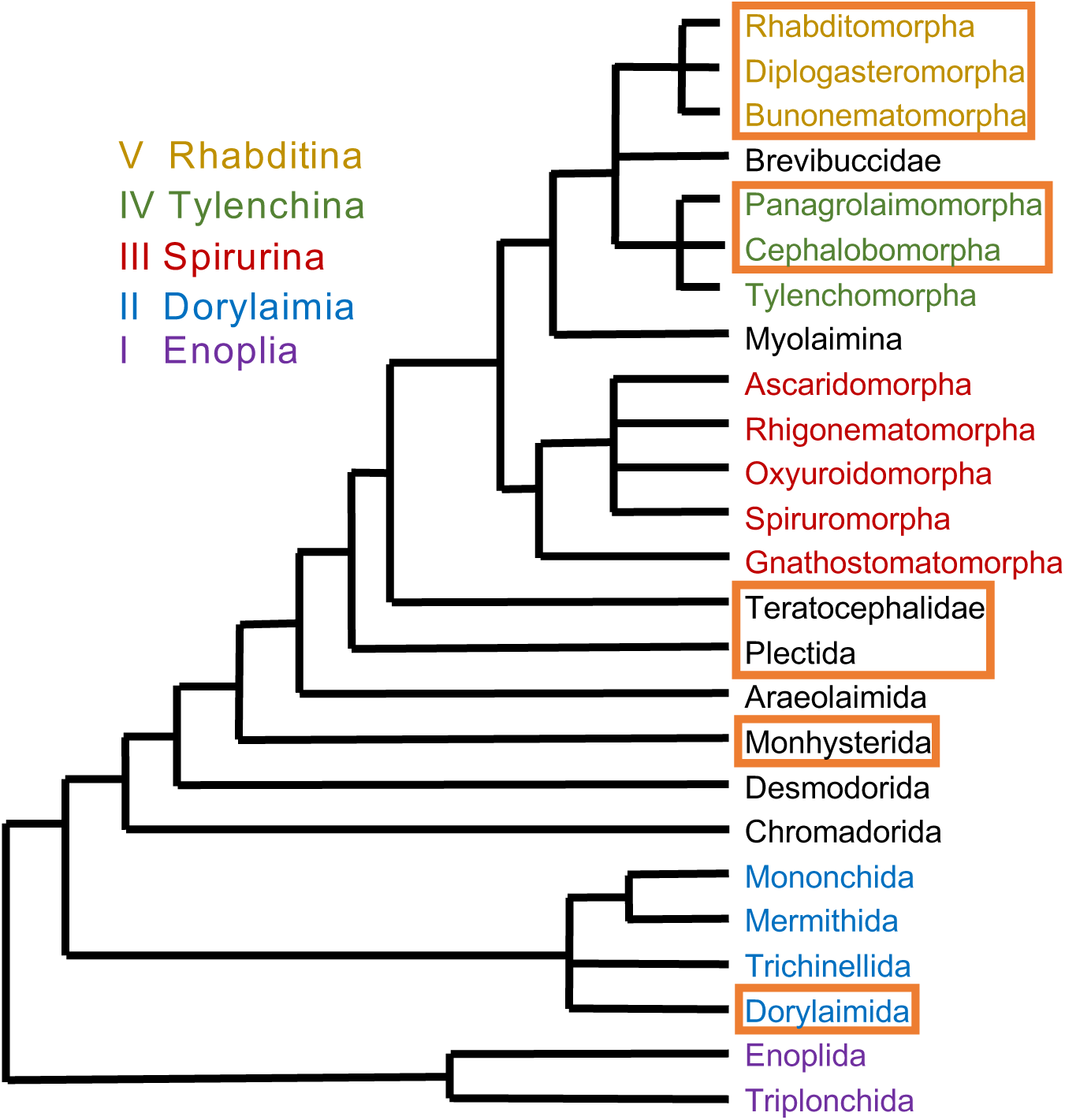
The nematode species isolated in our samples span several major clades of Nematoda. The phylogenetic tree of Nematode was redrawn based on (9) highlighting the spanned clades in this study.

**Figure S2.**
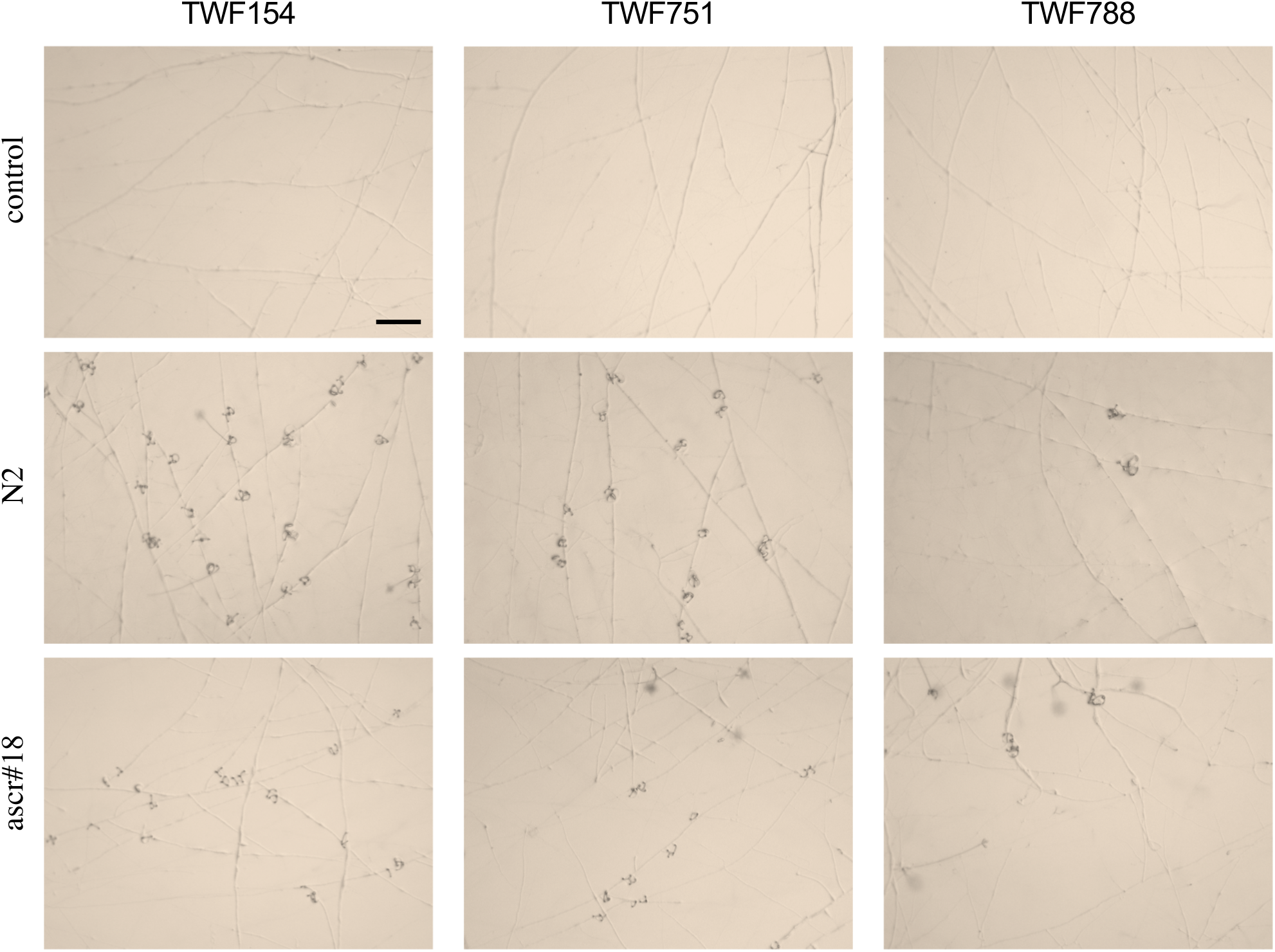
Images of the traps induced by *C. elegans* wild-type strain N2 or ascarosides (ascr#18) in *A. oligospora* wild isolates that were strongly (TWF154), intermediately (TWF751), or weakly (TWF788) responsive to the nematode cues. Bar = 200 µm.

**Figure S3.**
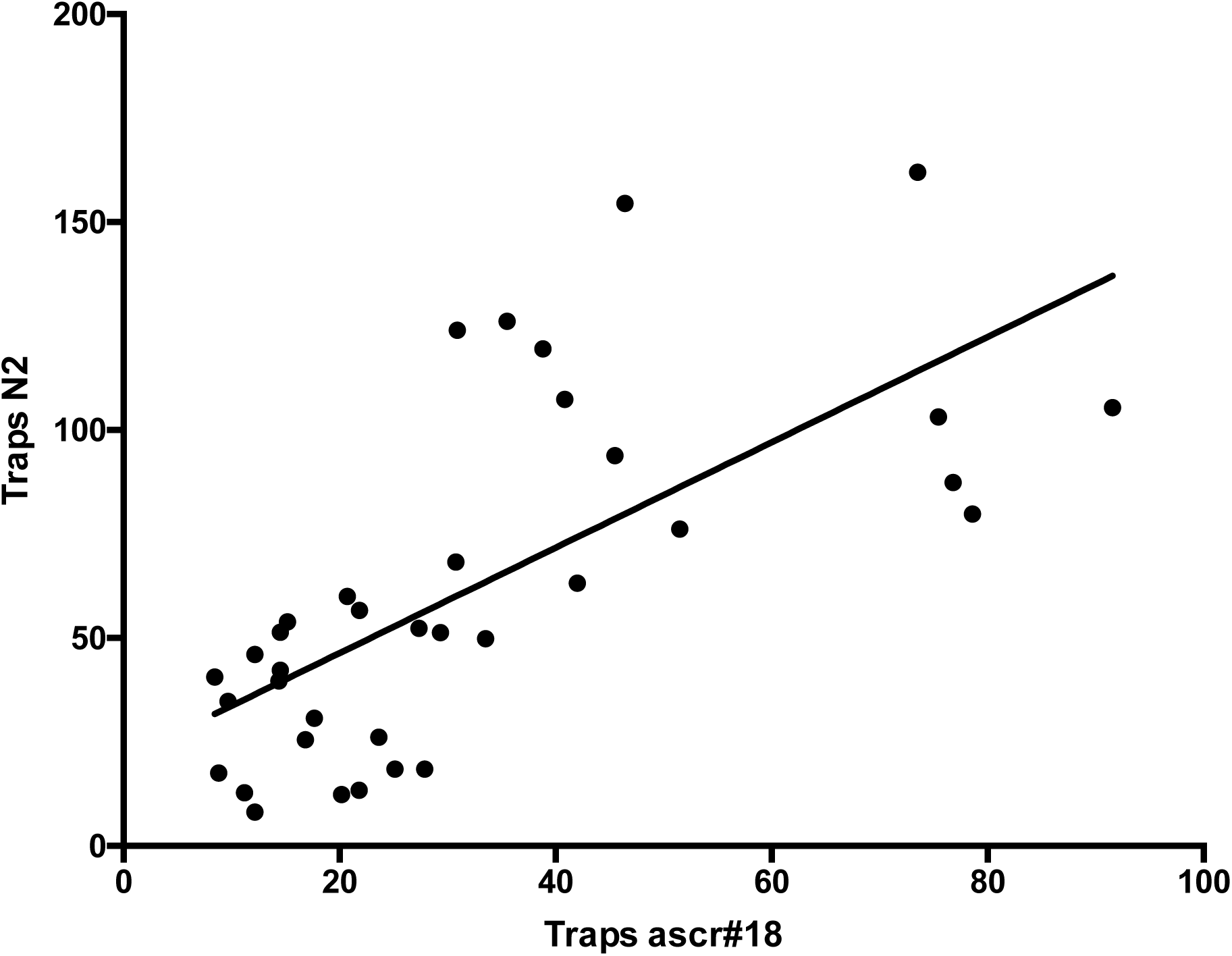
Correlation analysis of *A. oligospora* response to nematode versus ascarosides. Each dot represents a wild isolate of *A. oligospora*, the Y-axis represents the mean number of traps after exposition to *C. elegans* (N2) (Fig 1A) and the X-axis represents the mean number of traps in response to ascarosides (Fig1B). The line of best fit (Y = 1.267X + 21.02) obtained from linear regression (R^2^=0.4659) is also shown in the graph.

**Figure S4.**
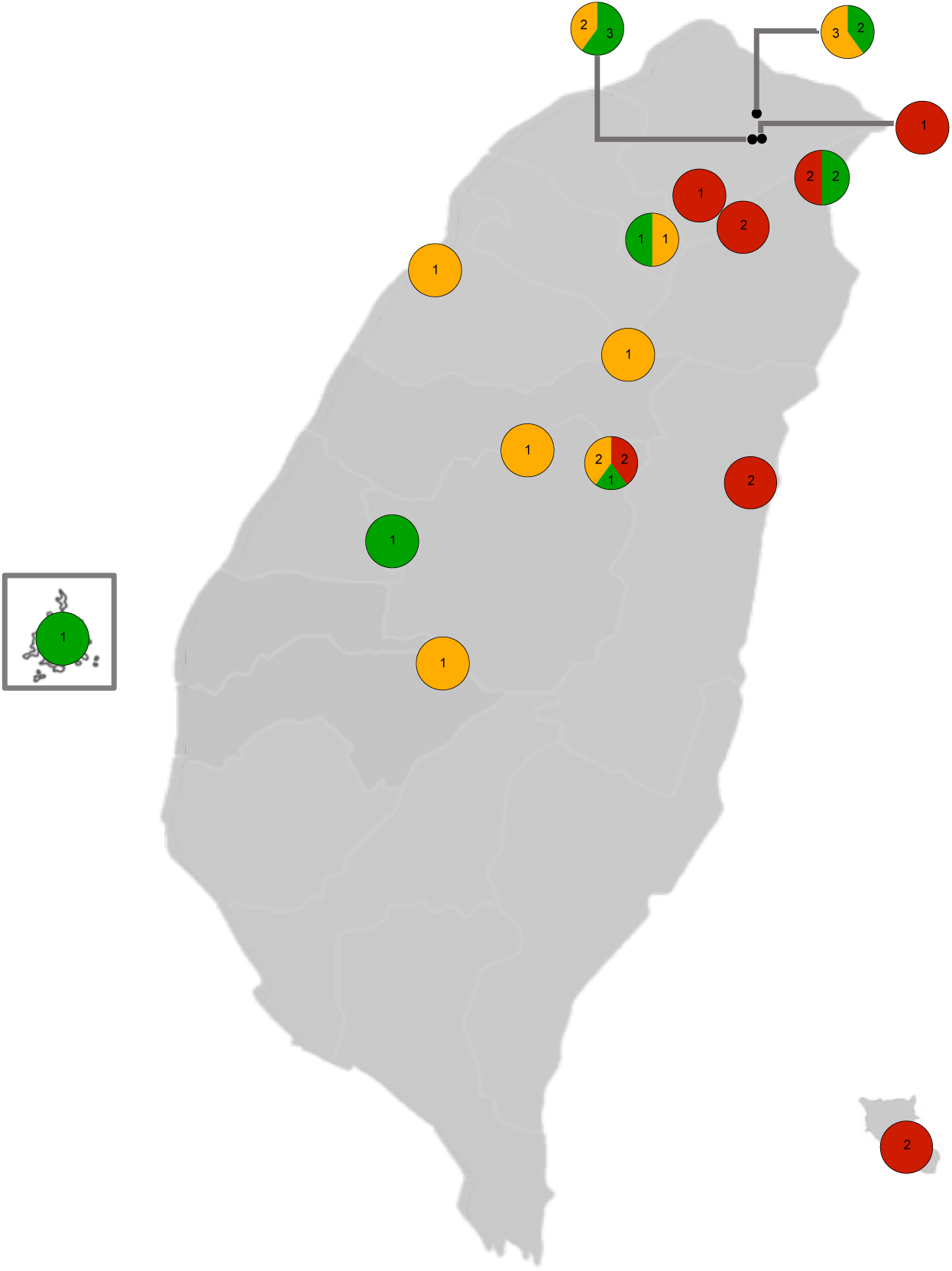
Geographical location of high, medium and low trapping-ability isolates of *A. oligospora*. Each pie chart shows the number of isolates exhibiting high (red), medium (yellow) and low (green) trapping ability at each sampling location. The trapping ability of the isolates was determined by their response to *C. elegans* (Fig 1A), isolates between TWF145-TWF899, TWF197-TWF162 and TWF703-TWF788 correspond to high, medium and low trapping ability, respectively.

**Figure S5.**
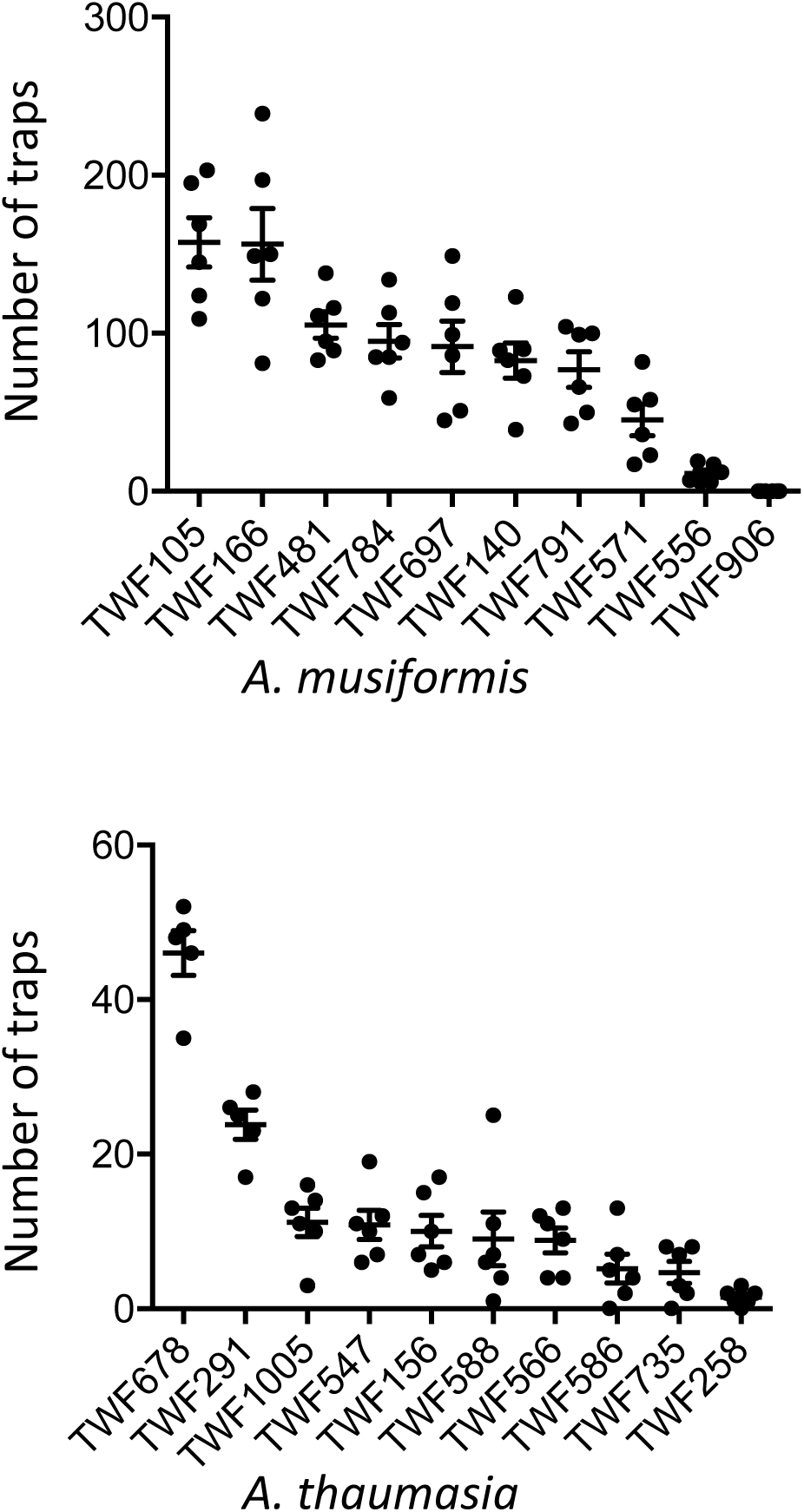
Prey-sensing ability variation among different isolates of the NTF *A. musiformis* and *A. thamausia.* Quantification of trap number induced by *C. elegans* wild-type strain N2 among wild isolates of *A. musiformis* (A) and *A. thamausia* (B) (mean ± SEM).

**Figure S6.**
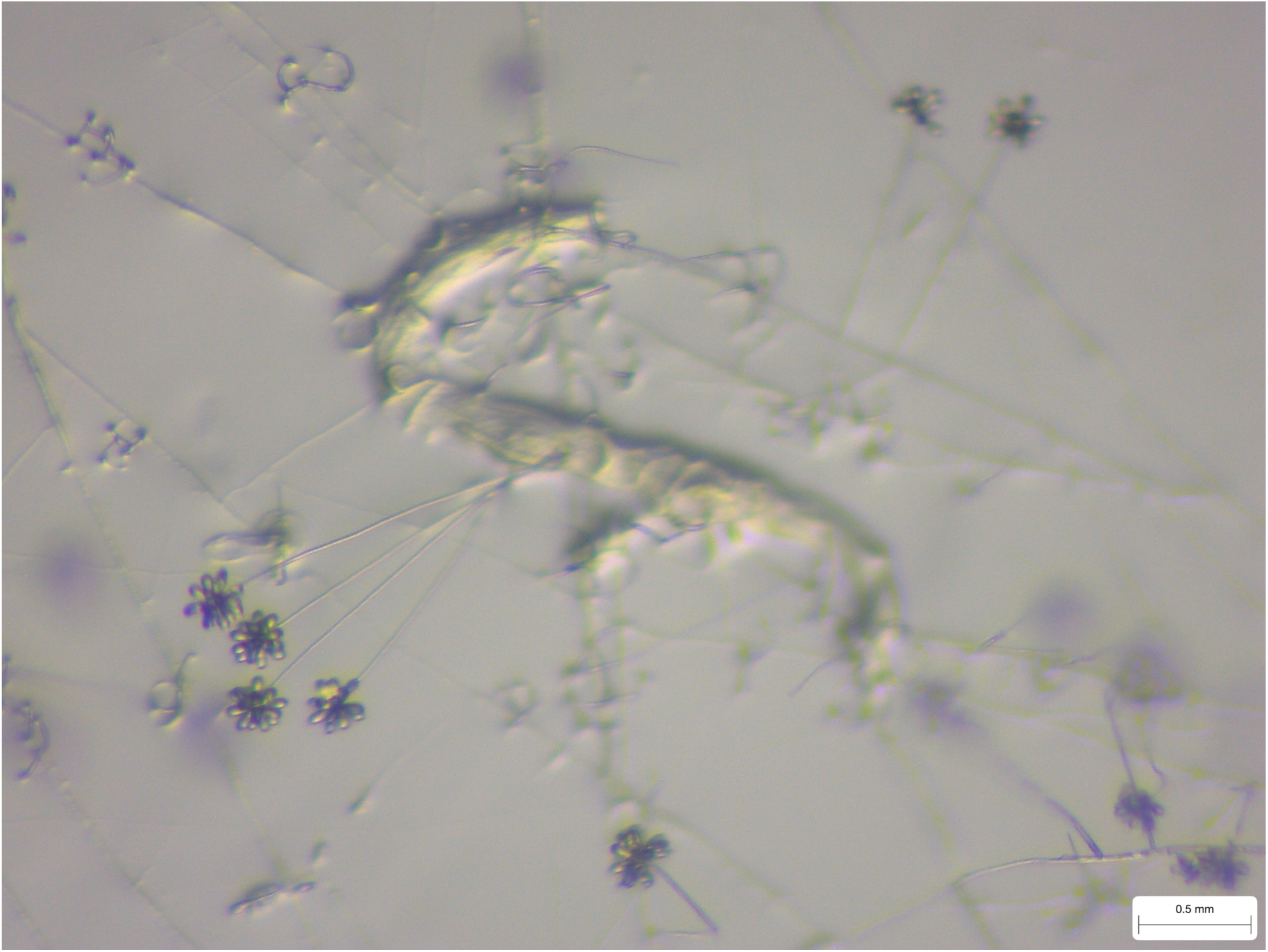
*A. oligospora* forms chains of conidia after consuming nematodes. The image depicts a *C. elegans* nematode trapped and consumed by *A. oligospora* (TWF154), and the conidia formed by the fungus after the prey consumption.

